# A unique adhesive motif of protein disulfide isomerase P5 supports its function via dimerization

**DOI:** 10.1101/2020.11.17.387910

**Authors:** Masaki Okumura, Shingo Kanemura, Motonori Matsusaki, Misaki Kinoshita, Tomohide Saio, Dai Ito, Chihiro Hirayama, Hiroyuki Kumeta, Mai Watabe, Yuta Amagai, Young-Ho Lee, Shuji Akiyama, Kenji Inaba

## Abstract

P5, also known as PDIA6, is a PDI-family member that plays an important role in the ER quality control. Herein, we revealed that P5 dimerizes via a unique adhesive motif contained in the N-terminal thioredoxin-like domain. This motif is apparently similar to, but radically different from conventional leucine-zipper motifs, in that the former includes a periodic repeat of leucine or valine residues at the third or fourth position spanning five helical turns on 15-residue anti-parallel α-helices, unlike the latter of which the leucine residues appear every two helical turns on ∼30-residue parallel α-helices at dimer interfaces. A monomeric P5 mutant with the impaired adhesive motif showed structural instability and local unfolding, and behaved as an aberrant protein that induces the ER stress response. Disassembly of P5 to monomers compromised its ability to inactivate IRE1α via reduction of intermolecular disulfide bonds and its Ca^2+^-dependent regulation of chaperone function *in vitro*. Thus, the leucine-valine adhesive motif supports structure and physiological function of P5.

## INTRODUCTION

The endoplasmic reticulum (ER) is an essential organelle where secretory and membrane proteins are newly synthesized and folded properly with the assist of numerous numbers of molecular chaperones and post-translational modification enzymes including disulfide bond formation catalysts(Bulleid & Ellgaard, 2011, Fass & Thorpe, 2018, Sevier & Kaiser, 2002). More than 20 members of the protein disulfide isomerase (PDI) family have been identified in mammalian cells, and they are believed to cooperate synergistically to promote oxidative folding of a wide variety of secretory proteins(Bulleid & Ellgaard, 2011, Okumura, Kadokura et al., 2015). Our previous *in vitro* studies demonstrated that among the PDI family members (PDIs), canonical PDI serves as a versatile enzyme that can selectively introduce native disulfide bonds into substrates, while P5 and ERp46, two other ubiquitously expressed PDIs(Cheng, Liu et al., 2017, Su, Cooke et al., 2002), are dedicated to rapid and promiscuous disulfide introduction during early oxidative folding(Kojima, Okumura et al., 2014, Sato, Kojima et al., 2013). Thus, PDI and P5/ERp46 likely work at different stages of oxidative folding, and act cooperatively to increase the production of multiple-disulfide proteins.

P5, also known as PDIA6, ERp5, TXNDC7, and CaBP1, comprises two redox-active thioredoxin (Trx)-like domains (**a^0^** and **a**) and a redox-inactive Trx-like domain **b** in this order from the N-terminus (Extended Data Fig. 1a)(Lundström-Ljung, Birnbach et al., 1995). Although crystal structures of isolated **a^0^** and **a** domains have been solved at atomic resolutions(Sato et al., 2013), the overall structure of P5 remains to be determined. In addition to functioning as a disulfide formation catalyst, proteomic analysis demonstrated that P5 forms a non-covalent complex with BiP(Jessop, Watkins et al., 2009), a representative ER-resident chaperone, suggesting that P5 and BiP may work cooperatively to maintain the protein homeostasis in the ER.

Although physiological functions of P5 are not fully understood, a previous study showed that P5 serves as a regulator of the unfolded protein response (UPR) by acting on the oligomers of inositol-requiring enzyme 1 α (IRE1α), one of the primary UPR sensors in human cells(Eletto, Chevet et al., 2014, Eletto, Eletto et al., 2016). Under ER-stressed conditions, IRE1α transiently oligomerizes in response to the accumulation of misfolded proteins, and intermolecular disulfide bonds are supposed to stabilize the oligomeric states(Eletto et al., 2014, Eletto et al., 2016, Matsusaki, Kanemura et al., 2019). When ER-stressed conditions are relaxed, the IRE1α oligomers must disassemble reversibly to dimers or monomers to terminate the UPR signaling. The detailed mechanism of this inactivation pathway remains unclear, but P5 was shown to cleave the intermolecular disulfide bonds in IRE1α oligomers to initiate the inactivation and negatively regulate the UPR(Amin-Wetzel, Saunders et al., 2017, Eletto et al., 2016). Additionally, it has been reported that P5 cleaves a specific disulfide bond buried inside the MHC class-I chain-related polypeptide A molecule on the surface of tumor cells and thereby promotes tumor immune evasion(Kaiser, Yim et al., 2007). Thus, several lines of evidence indicate that P5 plays essential roles in oxidative and reductive processes under physiological conditions.

In the present work, we extensively performed structural and biochemical analyses for human P5 and revealed its dimeric structure in which the N-terminal Trx-like **a^0^** domains of two protomers interact with each other via the unique Leu-Val adhesive motif. This motif is apparently similar to, but radically different from conventional Leu-zipper motifs in that the former contains a periodic repetition of Leu or Val residues at the third or fourth position spanning five helical turns on the 15-residue anti-parallel α-helices, unlike the latter of which the Leu residues appear every two helical turns on ∼30-residue parallel α-helices and constitute dimer interfaces. Upon the mutational impairment of this motif, P5 became monomeric and structurally destabilized in concomitant with local unfolding. Resultantly, the P5 mutant induced ER stress response when overexpressed in cells. The monomeric P5 also showed the compromised ability to inactivate IRE1α through reduction of intermolecular disulfide bonds and the diminished Ca^2+^-dependent regulation of chaperone function. Altogether, we revealed the essential roles of the Leu-Val adhesive motif for supporting structure and physiological functions of P5.

## RESULTS

### P5 dimerizes via the Leu-Val adhesive motif in the N-terminal Trx-like domain a^0^

To obtain information on both the overall shape and the domain arrangement of full-length P5 in solution, we carried out small-angle X-ray scattering (SAXS) analysis. Fig. 1a shows the SAXS profiles extrapolated to zero concentration for the reduced and oxidized forms of P5. Guinier plots were linear, without any upward curvature at low *Q*^2^ (Fig. 1a, inset), implying no aggregation for both redox forms. The values for radius of gyration, *R*g, were determined to be 47.70 ± 0.26 Å and 47.85 ± 0.28 Å for the reduced and oxidized forms, respectively, from the slope and intercept of linear fits of the Guinier plot (Table 1). P5 has a considerably larger radius than the U-shaped PDI molecules (36 Å for reduced and 37.5 Å for oxidized forms)(Okumura, Noi et al., 2019) and the open V-shaped ERp46 molecule (42 Å)(Kojima et al., 2014). Molecular mass, calculated from the normalized forward intensity, *I*(0), using bovine serum albumin (BSA; c.a. 66,400 Da) as the standard, was c.a. 97,700 Da for the reduced form and 101,500 Da for the oxidized form. Based on a calculated molecular mass of c.a. 48,400 Da in a monomer unit, both reduced and oxidized forms of P5 predominantly form dimers in solution (Table 1).

**Table 1.** SAXS structural parameters.

| | $R_g^a$ | $R_g^b$ | $I(0)^a$ | $I(0)^b$ | $D_{\max}^c$ | $MM^d$ | $MM^e$ | $MM^f$ | $MM^g$ | $MM^h$ | Oligomeric State |
| --- | --- | --- | --- | --- | --- | --- | --- | --- | --- | --- | --- |
|  | (Å) | (Å) | (a.u.) | (a.u.) | (Å) | (kDa) | (kDa) | (kDa) | (kDa) | (kDa) | - |
| P5-OX<sup>i</sup> | 47.70 ± 0.26 | 48.57 ± 0.33 | 39.14 ± 0.14 | 39.16 ± 0.15 | 174 | 97.7 | 117.0 | 97.6 | 109.1 | 48.4 | Dimer |
| P5-RED<sup>i</sup> | 47.85 ± 0.28 | 49.52 ± 0.35 | 40.66 ± 0.15 | 40.94 ± 0.17 | 180 | 101.5 | 115.7 | 95.3 | 107.5 | 48.4 | Dimer |
| P5-RED-5A<sup>i</sup> | 38.51 ± 0.28 | 40.08 ± 0.37 | 20.90 ± 0.10 | 20.92 ± 0.11 | 150 | 52.2 | 49.9 | 42.7 | 47.0 | 48.3 | Monomer |
| a0a-RED<sup>i</sup> | 37.01 ± 0.22 | 39.05 ± 0.18 | 28.12 ± 0.10 | 28.65 ± 0.10 | 134 | 70.2 | 70.3 | 68.2 | 72.6 | 32.1 | Dimer |
| ab-RED<sup>i</sup> | 31.88 ± 0.24 | 32.79 ± 0.27 | 16.59 ± 0.08 | 16.63 ± 0.08 | 120 | 41.4 | 45.8 | 42.8 | 40.9 | 35.1 | Monomer |
| BSA<sup>j</sup> | 27.84 ± 0.14 | n.d. | 26.61 ± 0.08 | n.d. | n.d. | n.d. | n.d. | n.d. | n.d. | 66.4 | Monomer |

<sup>a</sup> Guinier analysis using the $Q$ range from 0.01008498 Å<sup>-1</sup> (0.01004249 Å<sup>-1</sup> for P5-red-5A) to $Q_{\max} < 1.3 / R_g$ . <sup>b</sup> Estimates in real space upon $P(r)$ determination. <sup>c</sup> Maximum dimension estimated by using GNOM package. <sup>d</sup> Molecular mass calculated by using the $I(0)$ value for BSA as the standard. <sup>e</sup> Molecular mass estimated from <sup>e</sup> Porod volume(Petoukhov et al., 2012), <sup>f</sup> apparent volume(Fischer, de Oliveira Neto et al., 2009), and <sup>g</sup> volume of correlation(Rambo & Tainer, 2013) by using ATSAS suite(Franke, Petoukhov et al., 2017). <sup>h</sup> Theoretical molecular mass calculated according to amino acid sequences. <sup>i</sup> Extrapolated to infinite dilution. <sup>j</sup> At finite concentration of 1.77 mg/ml(Akiyama, 2010).

**Fig. 1.**
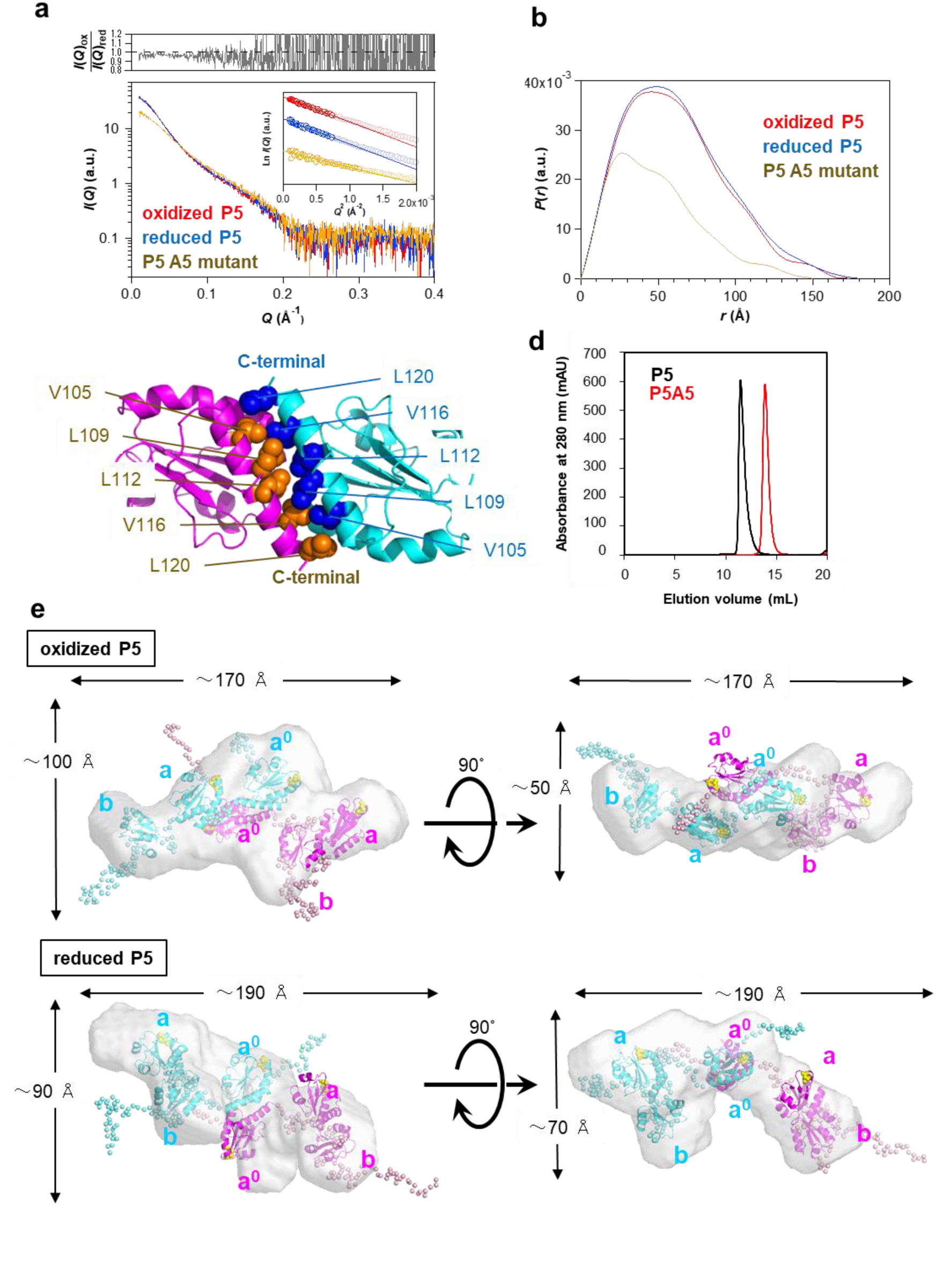
Overall structure of P5 and the role of the Leu-Val adhesive motif. **a,** SAXS profiles of oxidized P5 (red), reduced P5 (blue), and the P5A5 mutant (yellow). The inset shows Guinier plots generated using the *Q* range (highlighted data points in the inset) shown in Table 1. The upper panel shows an undulated deviation of *I*(*Q*)ox/*I*(*Q*)red from the unity value. **b,** Pair distribution function *P*(*r*) of oxidized P5 (red), reduced P5 (blue), and the P5A5 mutant (yellow). **c,** The crystal structure of P5 **a^0^** domain(Sato et al., 2013). Note that V105, L109, L112, V116, and L120 located on α-helix 4 in the **a^0^** domain interact tightly with each other at the interface. Note that V105, L109, L112, V116, and L120 are aligned on the same side of α-helix 4. **d,** Size-exclusion chromatography profiles of P5 (black) and P5A5 (red). **e,** Representative rigid-body refinement models of oxidized (upper) and reduced (lower) forms of P5 calculated using CORAL are represented by a ribbon diagram. Redox active sites in domains **a^0^** and **a** are shown by yellow spheres. (see Table 2 for more details).

To identify domains responsible for the dimerization, we prepared two sorts of truncated mutants, in which either the first Trx-like domain **a^0^** or the third Trx-like domain **b** (Extended Data Fig. 2a) including the C-terminal extension loop (residue 291-440) was deleted. Size-exclusion chromatography (SEC) analysis demonstrated that **a^0^**-**a** and **a**-**b** were eluted out at around the elution volumes of 12 and 15 ml, respectively (Extended Data Fig. 2b, c), suggesting that **a^0^**-**a** is significantly larger in size than **a**-**b**. Consistently, molecular masses estimated by SAXS analysis were c.a. 70,200 Da for **a^0^**-**a** and c.a. 41,400 Da for **a**-**b** based on the *I*(0) value from BSA as the standard (Extended Data Fig. 2d, e, and Table 1). Collectively, it is concluded that P5 forms a homodimer via the N-terminal **a^0^** domain.

To gain mechanistic insight into the dimerization via **a^0^**, we looked into our previously solved crystal structure of **a^0^** (Sato et al., 2013), and found that V105, L109, L112, V116, and L120 of α-helix 4 in **a^0^** form tight contacts with each other (Fig. 1c). It is widely known that a Leu zipper is a commonly used three-dimensional structural motif that promotes dimerization through adhesive force between two ‘parallel’ α-helices. This motif, in many cases, is a 30-amino acid segment and contains a periodic repetition of Leu residues at every seventh position spanning eight helical turns(Landschulz, Johnson et al., 1988). However, the adhesive motif found in P5 **a^0^** consists of a 15-amino acid segment with a periodic repetition of Leu or Val residues at the third or fourth position spanning five helical turns, and these residues are located on ‘anti-parallel’ α-helices. BLAST analysis revealed that this motif is highly conserved among the orthologues of P5 (PDIA6) (Extended Data Fig. 2f). To confirm that V105, L109, L112, V116, and L120 play an essential role in P5 dimerization in solution, we mutated these residues to Ala and analyzed the oligomeric state of the resultant mutant (named P5A5 in this paper). The elution volume was shifted from 11.5 mL for P5 to 15 mL for P5A5 (Fig. 1d), and SAXS analysis clearly showed that P5A5 had a molecular weight corresponding to the monomer (Fig. 1a, b, and Table 1). We thus verified that P5 forms a homodimer via the unique adhesive motif in the N-terminal **a^0^** domain.

### Overall structures of P5

To determine the molecular shape of P5, the pair distribution function, *P*(*r*), was calculated from the observed SAXS curves using the GNOM package (Svergun, 1991). *P*(r) of reduced P5 was highly similar to that of oxidized P5, indicating that the overall structure of P5 is unaffected by the redox state (Fig. 1b). The largest linear distance values (*D*max) were estimated to be 174 Å for the oxidized form and 180 Å for the reduced form (Table 1). These values were significantly larger than those of PDI (126 and 138 Å for reduced and oxidized states, respectively) ^25^ and ERp46 (137 and 141 Å for reduced and oxidized states, respectively)(Kojima et al., 2014), consistent with the dimeric structure of P5. To draw the overall structures of P5, we performed rigid body modelling based on the present SAXS data, using the previous crystallographic data of P5 **a^0^** (PDB code: 3VWW)(Sato et al., 2013), P5 **a** (PDB code: 4GWR) and PDI **b** (PDB code: 4EKZ)(Wang, Li et al., 2013) and the program CORAL(Petoukhov, Franke et al., 2012) (Table 2). As the consequence of the rigid body refinement, multiple structure models with different Trx-like domain arrangements were generated as feasible conformations with reasonable fitness values (χ value of lower than 1.325 for oxidized form and 1.394 for reduced form; Table 2 and Extended Data Fig. 3). In the representative model judged with the lowest NSD value from DAMAVER package (Table 2), the six Trx-like domains contained in a P5 dimer are placed in a plane whose dimensions are ∼170 Å × ∼110 Å for the oxidized form and ∼190 Å × ∼110 Å for the reduced form (Fig. 1e). Except for the **a^0^** domains that are tightly packed via the adhesive motif, all domains are located far apart from each other, with very few physical contacts between them, resulting in various extended conformations of P5 (Extended Data Fig. 3). Such loosely-packed domain arrangement suggests a flexible nature for P5. The redox-active CXXC motifs of domains **a^0^** and **a** are both solvent-exposed and separated from each other (Fig. 1e), as they are in ERp46(Kojima et al., 2014).

**Table 2.** SAXS shape-reconstruction statistics for oxidized- and reduced-P5.

|  | **Oxidized P5** | **Reduced P5** |
| --- | --- | --- |
| Shape Reconstruction | CORAL | CORAL |
| $Q$ range ($\text{\AA}^{-1}$) | 0.01008 – 0.17462 | 0.01008 – 0.17462 |
| Symmetry | P1 | P1 |
| Total Number of Residues | 442 | 442 |
| Total Number of Dummy Residues | 129 | 129 |
| Dummy Residues | 1 – 26, 143 – 162, 275 – 278, 305 – 308<br>322 – 330, 339 – 344, 364 – 371, 391 – 442 | 1 – 26, 143 – 162, 275 – 278, 305 – 308<br>322 – 330, 339 – 344, 364 – 371, 391 – 442 |
| Total Number of Known Residues | 313 | 313 |
| Known Residues Treated as Rigid Bodies | 27 – 142, 163 – 274, 279 – 304, 309 – 321, 331 – 338, 345 – 363, 372 – 390 | 27 – 142, 163 – 274, 279 – 304, 309 – 321, 331 – 338, 345 – 363, 372 – 390 |
| SQRT($\chi^2$) (mean ± S.D.) | 1.178 – 1.325 (1.219 ± 0.042) | 1.122 – 1.394 (1.202 ± 0.079) |
| Number of Models Reconstructed | 20 | 20 |
| DAMAVAR NSD (mean ± S.D.)<sup>a</sup> | 3.891 – 4.760 (4.230 ± 0.240) | 4.344 – 5.310 (4.670 ± 0.217) |

<sup>a</sup>NSD: normalized spatial discrepancy

### The C-terminal Asp/Glu-rich segment of P5 is responsible for Ca^2+^ binding

In the preceding work, P5 was shown to bind calcium ions (Ca^2+^) and is hence also called CaBP1^7^. We herein analyzed binding of Ca^2+^ to P5 by isothermal titration calorimetry (ITC) (Fig.2b & Table 3). A series of ITC thermograms revealed Ca^2+^ binding ability of P5, depending on its constructs. P5 and P5A5 showed endothermic ITC peaks, indicating that P5 binds Ca^2+^ irrespective of the adhesive motif (Fig. 2b, upper panels). The ability of P5 to bind Ca^2+^ was retained after the deletion of the N-terminal **a^0^** domain or **a^0^-a** domains; P5-**ab** and P5-**b** displayed endothermic ITC peaks, like P5 (Fig. 2b, upper panels). By contrast, P5 did not exhibit appreciable reaction heat after the deletion of domain **b** and the subsequent C-terminal segment (Fig. 2b, lower left). Similarly, P5 with the deletion of residue 422-440 (ΔC) or the mutations of Asp/Glu to Asn/Gln (NQ) in the C-terminal segment did not show marked ITC peaks (Fig. 2b, lower middle and right). Thus, Ca^2+^ binding to P5-**a^0^a**, P5-ΔC and P5-NQ was negligible, not allowing reliable ITC analyses.

**Fig. 2.**
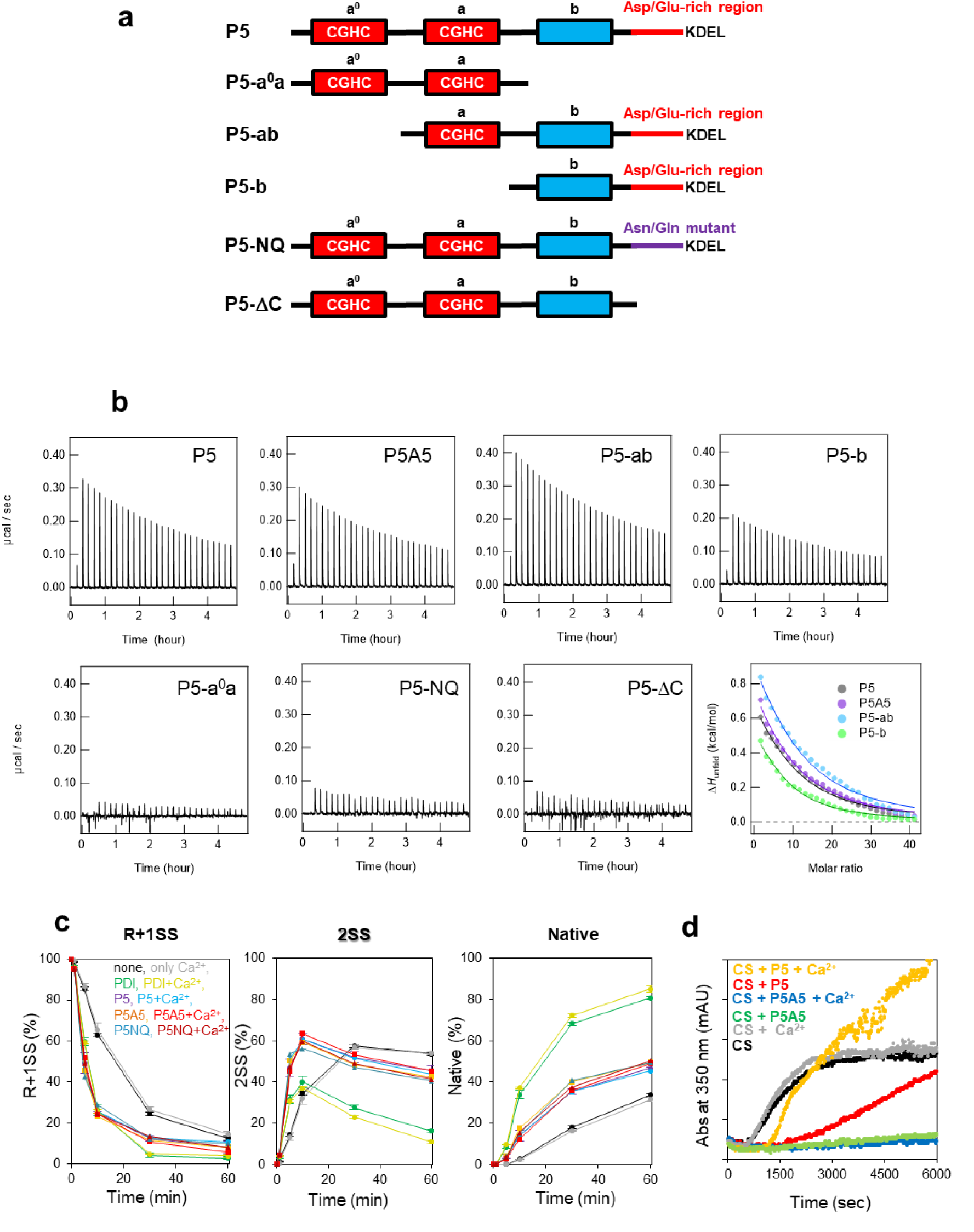
Ca^2+^ binding to P5 via the C-terminal Glu/Asp-rich segment. **a,** Domain organization of P5 and its mutants used in this study. Redox-active and redox-inactive Trx-like domains are indicated by red and blue boxes, respectively. Active-site sequences in the redox-active domains are indicated in the red boxes. **b,** ITC data (upper and lower left three panels) and binding isotherm data (lower right panel) for titration of Ca^2+^ against P5 and its mutants. Thermodynamic parameters for Ca^2+^ binding to P5 and its mutants are compiled in Table 3. The experiment was independently repeated three times with reproducible results. **c,** Quantitative analysis of oxidative folding of BPTI (30 μM) catalyzed by PDI, P5, and its mutants (each 1 μM) in the presence of GSH/GSSG (1 mM/0.2 mM). R, 1SS, 2SS, and N indicate the reduced, one disulfide-bonded, two disulfide-bonded, and native species, respectively. The occupancies of R, 1SS, 2SS, and N species are plotted as a function of folding time. Values are means ± SD of three independent experiments. **d,** Chaperone activity of P5 and the P5A5 mutant (8 μM) in the presence or absence of 1 mM CaCl2. Chaperone activity was assessed by monitoring absorbance at 350 nm with the use of citrate synthase (1 μM) as a model substrate. The same trend was observed in all three independent experiments.

**Table 3.** Thermodynamic parameters for Ca^2+^ binding to P5 and its mutants.

| | *n* | *K<sub>d</sub>*<br>(mM) | $\Delta H_{\text{bind}}$<br>(J/mol) | $\Delta G_{\text{bind}}$<br>(J/mol) | $-T\Delta S_{\text{bind}}$<br>(J/mol) |
| --- | --- | --- | --- | --- | --- |
| **P5** | 7.3±1.7 | 0.5±0.1 | 38.4±10.5 | -1,064.0±113.0 | -1,102.4±113.5 |
| **P5A5** | 6.7±1.3 | 0.6±0.1 | 45.5±10.2 | -1,061.7±84.4 | -1,107.2±85.0 |
| **P5-ab** | 8.2±1.9 | 0.6±0.1 | 51.2±14.3 | -1,050.3±119.9 | -1,101.5±120.7 |
| **P5-b** | 6.5±1.1 | 0.4±0.1 | 24.9±4.8 | -1,119.8±95.9 | -1,144.7±96.0 |
| **P5-NQ** | N.D. | N.D. | N.D. | N.D. | N.D. |
| **P5-ΔC** | N.D. | N.D. | N.D. | N.D. | N.D. |
| **P5-a<sup>0</sup>a** | N.D. | N.D. | N.D. | N.D. | N.D. |

Thermodynamic analyses provided quantitative information on the P5-Ca^2+^ complex formation (Fig. 2b, lower right panel). Affinity for Ca^2+^ of P5 was weak in nature, and, not largely different among P5, P5A5, P5-**ab**, and P5-**b** on the basis of the changes in Gibbs free energy (Δ*G*bind) of ca. -1.1 kcal/mol and the dissociation constant (*K*d) ranging from 0.4 - 0.6 mM (Table 3). Taking into consideration of the ER Ca^2+^ concentration of 0.5 – 1 mM(Meldolesi & Pozzan, 1998), however, this observation suggests the feasibility of Ca^2+^ binding to P5 under physiological settings. Binding stoichiometric *n* values indicate that 6 to 8 Ca^2+^ ions bind to each molecule of P5, P5A5, P5-**ab**, and P5-**b**. Intriguingly, Ca^2+^ binding to these three P5 constructs was purely driven by the favorable entropy change (Δ*S*bind) (i.e., Δ*S*bind > 0) (Table 3), which might be ascribed to complexation-induced dehydration that compensates for the energetically unfavorable enthalpy change (i.e., Δ*H*bind > 0).

The observations that P5A5, P5-**ab**, and P5-**b** retained 6 to 8 Ca^2+^-binding sites with similar affinity to P5 suggest that the dimerization is unlikely to affect the Ca^2+^-binding ability of P5. Moreover, Ca^2+^ binding to P5-**ab**, but not to P5 **a^0^a** and P5-ΔC, suggest that Ca^2+^ binding sites of P5 are located in the C-terminal segment. In line with this, the C-terminal segment of P5 contains negatively-charged residues (Asp and Glu) highly conserved across different species (Extended Data Fig 4). Upon the deletion of the mutations of Asp/Glu to Asn/Gln (NQ) in this segment, P5 was unable to bind Ca^2+^ (Fig. 2b). Collectively, we conclude that P5 binds 6 to 8 Ca^2+^ via the Asp/Glu-rich segment near the C-terminus.

### P5 catalyzes oxidative folding, irrespective of dimerization

We next sought to explore the role of dimerization in catalyzing oxidative folding. Oxidative folding assays were carried out using bovine pancreas trypsin inhibitor (BPTI) as a model substrate, of which the folding pathways have been characterized in detail *in vitro*(Okumura et al., 2019, Weissman & Kim, 1991) (Extended Data Fig. 5a). The reversed-phase high-performance liquid chromatography (RP-HPLC) profiles demonstrated that P5 under the redox conditions employed (GSH/GSSG = 1 mM/0.2 mM)(Walker, Lyles et al., 1996) converted the fully reduced form (R) to the partially or fully oxidized forms within 5 min, while PDI took 10 min until R completely disappeared (Extended Data Fig. 5b). P5 displayed numerous elution peaks between on-pathway 2SS N* des[30-51] and R at the refolding times of 1 min and 5 min, indicating that various off-pathway 1SS products accumulated during the early stage of P5-catalyzed oxidative folding. Although PDI diminished R species more slowly than P5, PDI generated larger amount of native-state BPTI within a 60 min folding time (Extended Data Fig. 5b). This result suggests that PDI converts folding intermediates of BPTI to a native species more efficiently than P5. Monomeric P5A5 gave almost the same results as P5 (Fig. 2c and Extended Data Fig. 5), indicating that dimerization does not affect the ability of P5 to catalyze BPTI oxidative folding.

### P5 chaperone function is regulated by Ca^2+^

To investigate how Ca^2+^ binding influences P5 function, we next carried out oxidative folding assays in the presence of Ca^2+^. During both the initial and subsequent steps of BPTI folding, P5, P5A5, P5 NQ, and PDI gave almost the same HPLC profiles with and without Ca^2+^. Thus, Ca^2+^ is not a regulator of the catalytic activity of P5 and PDI for promoting oxidative folding (Fig. 2c and Extended Data Fig. 5c).

Regarding the relationship between the chaperone function and Ca^2+^ binding, a previous study demonstrated that PDI induces client aggregation in the presence of Ca^2+^ (Primm, Walker et al., 1996), indicating Ca^2+^-dependent anti-chaperone function of PDI. We here investigated the effect of Ca^2+^ on P5 chaperone activity using citrate synthase (CS) as a model substrate having no disulfide bonds(Buchner, Grallert et al., 1998). Thermally aggregated CS was monitored by measuring absorbance at 350 nm (Fig. 2d, black dots). We thereby observed that increase of absorbance at 350 nm was significantly suppressed in the presence of P5, indicating its chaperone activity. Notably, P5A5 inhibited CS aggregation even more greatly than P5, which suggests that the dimerization negatively regulates P5 chaperone function. It is also to be noted that the addition of Ca^2+^ significantly diminished the chaperone activity of P5 (Fig. 2d, yellow and red dots, respectively), although thermal aggregation of CS in the presence of P5A5 was barely affected by Ca^2+^ (Fig. 2d, blue and green dots, respectively). Thus, P5 chaperone function is significantly regulated by Ca^2+^, and this regulation is dependent on its dimeric conformation.

### Monomeric P5 hampers reduction of an SS-linked IRE1**α** oligomer

A previous study demonstrated that P5 reduces disulfide-linked oligomeric species of IRE1α in cultured cells, and thereby facilitates the inactivation of IRE1α signaling(Eletto et al., 2016). To examine whether P5 directly cleaves the intermolecular disulfide bonds of IRE1α *in vitro* (Fig. 3), we prepared the recombinant disulfide (SS)-linked IRE1α luminal domain (LD), which consists of two monomers linked by an intermolecular disulfide bond. SS-linked IRE1α LD was purified homogeneously following the procedures illustrated in Extended Data Fig. 6. Using the SS-linked IRE1α LD, we performed gel-shift assays to probe the intermolecular disulfide cleavage by P5 in the presence of TCEP, an irreversible reducing agent (Fig. 3a). In the absence of PDI family enzymes (no PDIs), the SS-linked IRE1α LD (IRE1SS) decreased only slightly in concomitant with slow generation of the cleaved IRE1α LD (IRE1SH) during the incubation time (Fig. 3b, bottom). Notably, the presence of P5 greatly accelerated the decay of the IRE1SS species, and the IRE1SH species increased much more rapidly than when no PDIs were added (Fig. 3b, top). Quantification of the band intensity confirmed that P5 decreased IRE1SS to 16% after 30-min incubation (Fig. 3c, upper panel). The rate of P5-mediated IRE1SH species generation was not altered in the presence of 1 mM CaCl2 (Fig. 3e), although the chaperone function of P5 was regulated by Ca^2+^ (Fig. 2d). Thus, the IRE1α reduction activity of P5 is insensitive to Ca^2+^.

**Fig. 3.**
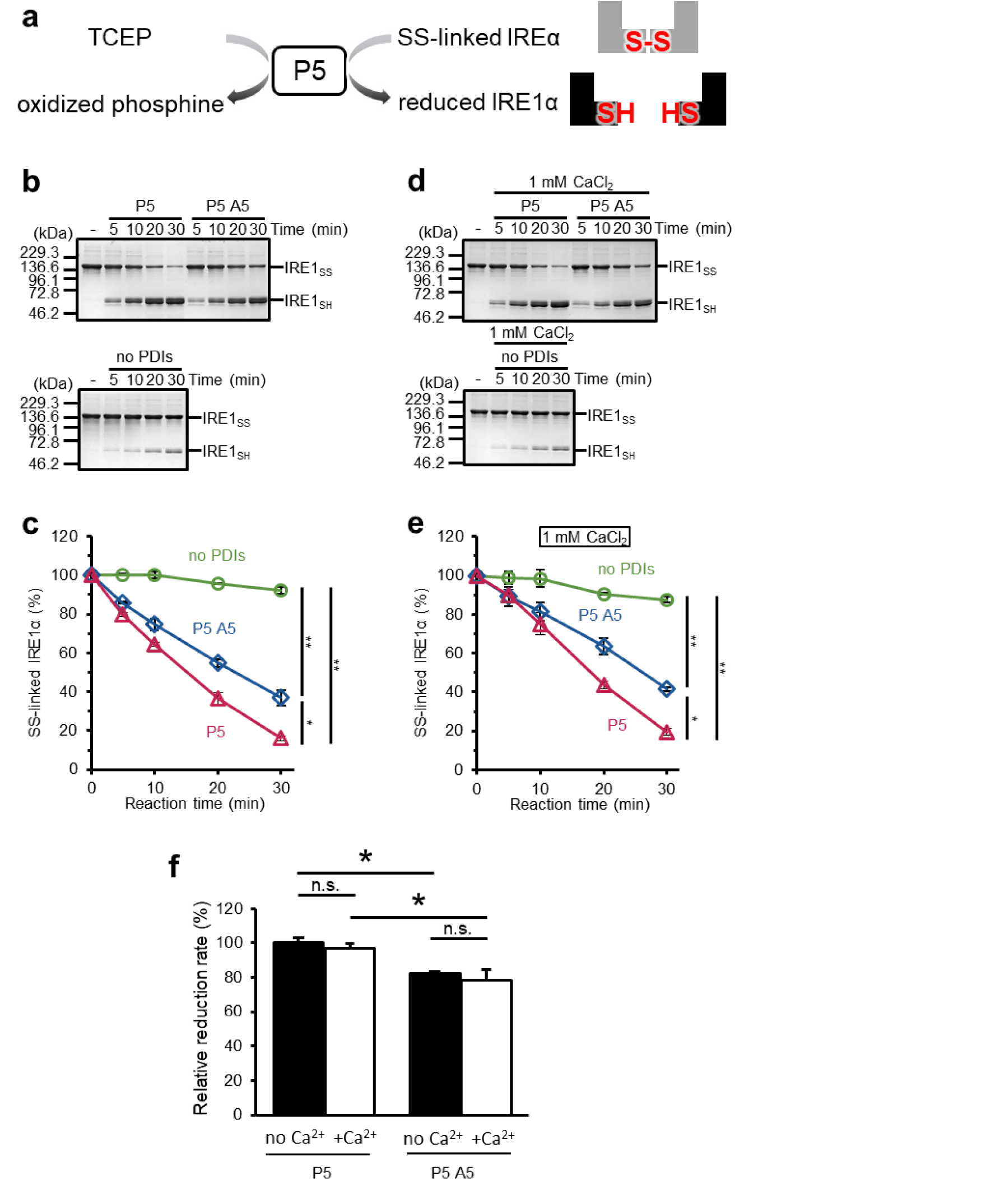
Role of the Leu-Val adhesive motif in P5-mediated disulfide bond reduction. **a,** Schematic representation of the disulfide-bonded IRE1α reduction assay. Disulfide-bonded species of the IRE1α (SS-linked IRE1α) luminal domain (LD) are reduced by P5 using tris(2-carboxyethyl)phosphine (TCEP) as a reducing source. **b**, Time course of P5-mediated reduction of the SS-linked IRE1α LD. Purified and SS-linked IRE1α LD (25 µM) was incubated without P5 as a control, with 0.5 μM P5 or P5A5 in the presence of 250 µM TCEP. The reactions were quenched with N-ethylmaleimide (NEM) at indicated time points. Reaction mixtures were separated by non-reducing 7.5% SDS-PAGE. IRE1SS and IRE1SH represent SS-linked IRE1α dimers and IRE1α monomers, respectively. **c,** Quantitative analysis of the SS-linked IRE1α dimer species shown in panel b. Values represent means ± standard error of the mean (SEM) from three independent experiments. Statistical analysis of the relative intensities of the SS-linked IRE1α dimer species at 30 min of reaction time was performed using one-way analysis of variance (ANOVA) and Tukey-Kramer tests. **d,** Results of the same experiments described in panel b but with 1 mM CaCl2. **e**, Quantitative analysis of the SS-linked IRE1α dimer species shown in panel d. Values are means ± SEM from three independent experiments. Statistical analysis of the relative intensities of the SS-linked IRE1α dimer species at 30 min of reaction time was performed using one-way ANOVA and Tukey-Kramer tests. **f,** Relative insulin reduction rate for P5 in the absence (black bar) and presence (white bar) of 1 mM CaCl2. Relative insulin reduction rate was calculated from NADPH consumption with or without 1 mM CaCl2, in the presence of 30 μM insulin, 10 mM GSH, 200 μM NADPH, 2 U/mL glutathione reductase, and 1 µM P5 or P5A5. Data are represented as means ± SD (n = 3; n.s., not significant; \**p* <0.05; \*\**p* <0.01; Tukey-Kramer test).

Of note, the mutations at the P5 dimerization motif significantly compromised the intermolecular disulfide cleavage activity of P5; P5A5 decreased the IRE1SS species to 37% after 30-min incubation time (Fig. 3b and 3c). Thus, the dimeric structure of P5 serves to promote dissociation and inactivation of IRE1α via disulfide reduction.

To further examine the higher disulfide reduction activity of P5 than its monomeric mutant, we performed insulin reduction assays(Morjana & Gilbert, 1991). The results showed that P5A5 possessed lower insulin reduction activity than P5, and this was barely affected by the presence of Ca^2+^ (Fig. 3f and Extended Data Fig. 7). Thus, Ca^2+^ effect was observed only marginally in disulfide reduction by P5, although dimerization *per se* plays a significant role in its reductive activity.

### Different impacts of overexpression of P5 and its monomeric mutant on cells

To explore the physiological significance of the P5 dimer, we investigated the effects of transient expression of C-terminal 3×Myc-tagged P5 in cultured cells (Fig. 4). After optimization of transfection conditions, P5 and the P5A5 mutant were transiently expressed at almost the same level in HeLa Kyoto cells (Fig. 4, left middle), and both co-localized with PDI, an ER marker (Extended Data Fig. 8). The impact of P5 overexpression on cells was assessed by monitoring the expression levels of BiP and GRP94, representative ER stress markers(Kozutsumi, Segal et al., 1988). Notably, in contrast to that of P5, expression of P5A5 caused a significant increase in the expression levels of BiP and GRP94 (Fig. 4a, left top and right). Thus, expression of monomeric P5 appears to somehow affect the ER homeostasis. Considering that IRE1 activation can induce a modest upregulation of BiP under ER-stressed conditions(Lee, Iwakoshi et al., 2003), the expression of P5A5 with the compromised IRE1 disulfide reduction activity (Fig. 3) may have led to the higher IRE1 activation. Alternatively, overexpressed P5A5 may have behaved as an aberrant protein that induces the ER stress, resulting in the upregulation of BiP and GRP94.

**Fig. 4.**
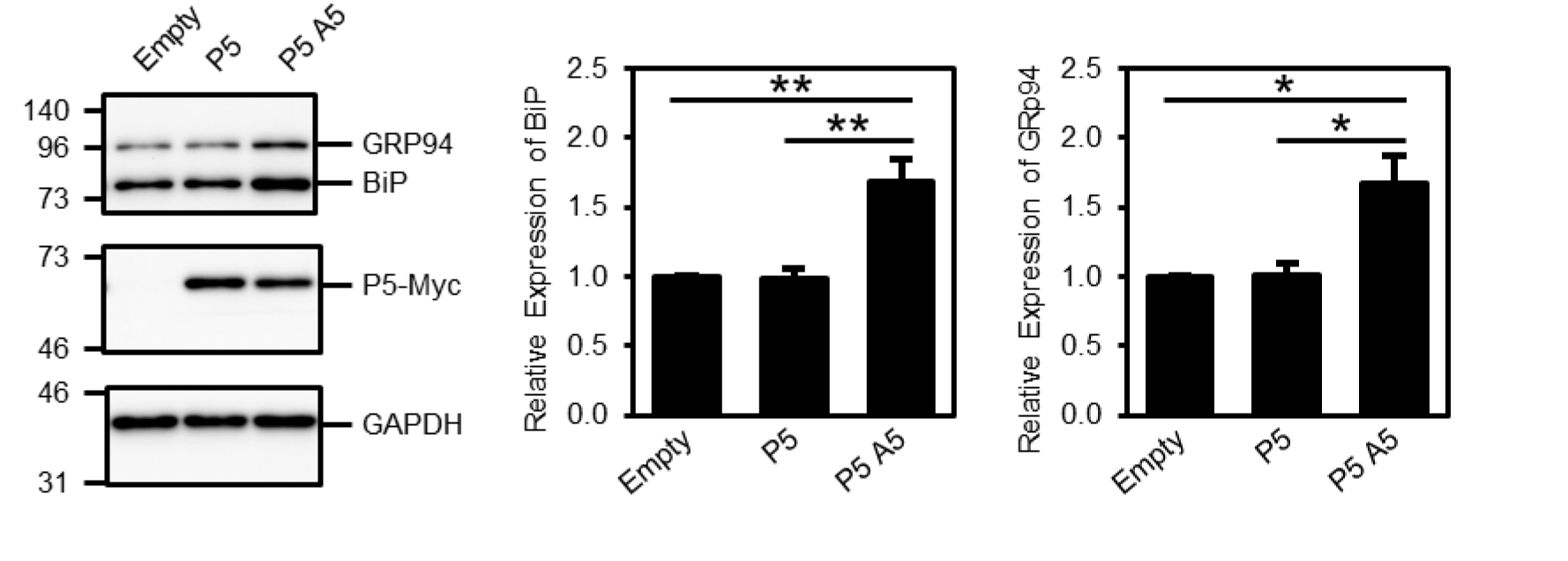
Physiological significance of P5 dimers in cells. Induction of BiP expression by overexpression of P5 mutants. (Left) HeLa Kyoto cells transfected with vector control (Empty) or P5-Myc-KDEL (P5, P5A5) were incubated for 40 h. Cells were harvested in SDS sample buffer containing N-ethylmaleimide (NEM) and analyzed by immunoblotting with anti-KDEL (top), anti-cMyc (middle), and anti-glyceraldehyde-3-phosphate dehydrogenase (GAPDH; bottom) antibodies. (Center, Right) The signal intensity of BiP or GRP94 in P5- or P5A5-overexpressing cells relative to that of BiP or GRP94 in Empty vector cells was quantified. Data are presented as means ± SD (n = 3;; \**p* <0.05; \*\**p* <0.01; Tukey-Kramer test).

### Loss of P5 dimerization induces conformational destabilization

To explore the possibility that overexpressed monomeric P5 serves as an aberrant protein that can induce the ER stress, we extensively investigated structure and physicochemical properties of P5A5 and compared them with those of P5. To this end, the thermal denaturation analysis was performed for P5 and P5A5 by measuring far-UV circular dichroism (CD) spectra at different temperatures (Fig. 5a). Thermodynamic parameters of heat denaturation were estimated based on the two-state transition. Of note, P5 displayed the midpoint temperature for denaturation (*T*m) of 55.3 °C, which was higher by 6.2 °C than that of P5A5 (49.1 °C) (Table 4). The change in enthalpy of unfolding (Δ*H*unfold) was calculated to be ∼590 and ∼230 kJ/mol for P5 and P5A5, respectively (Table 4). These results suggest that the dimerization contributes in the overall conformational stability.

**Fig. 5.**
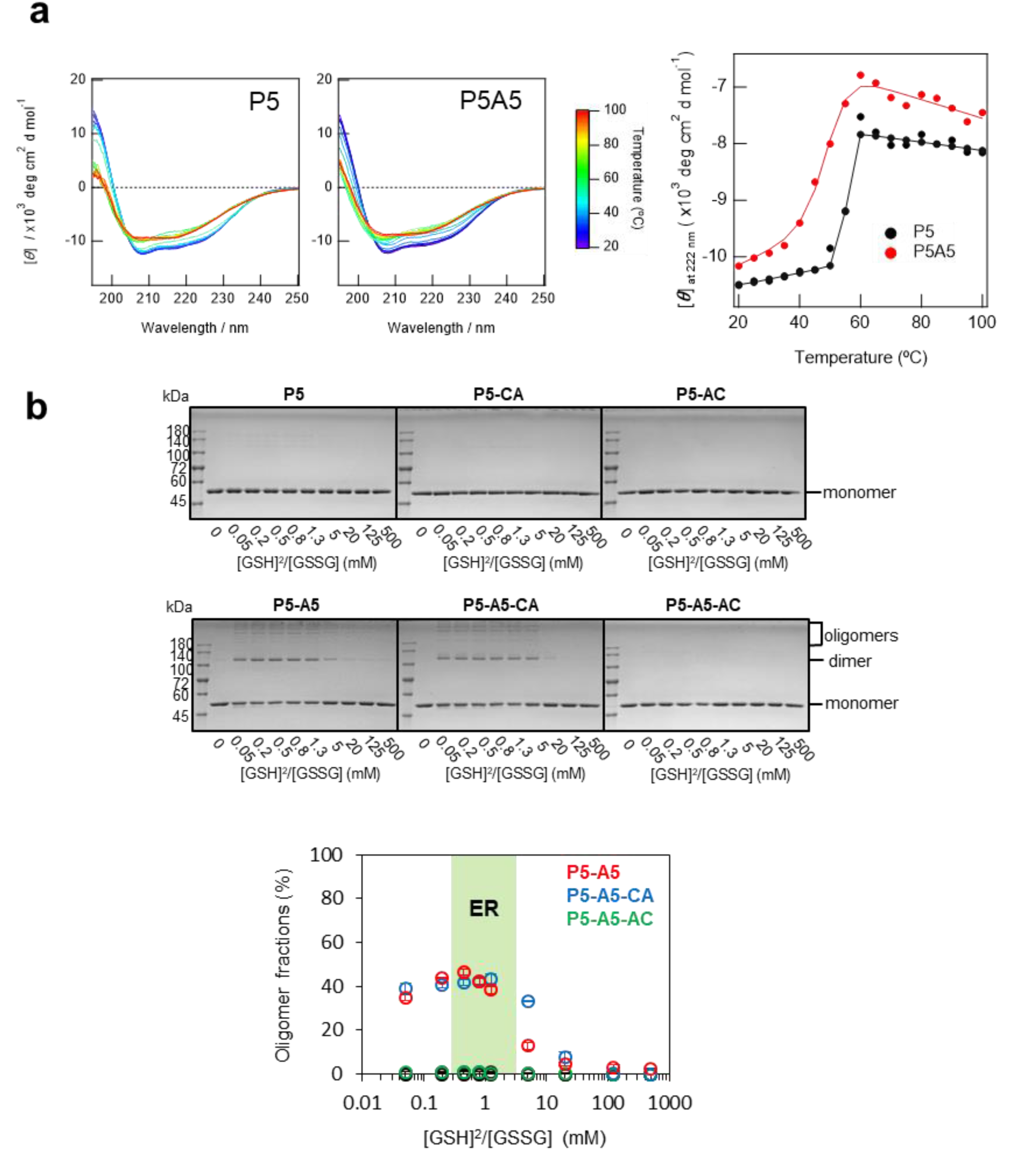
Loss of the Leu-Val adhesive motif causes thermodynamic destabilization and artefactual disulfide-linked oligomerization. **a,** Heat denaturation of P5 or P5A5 thermodynamically analyzed by using the CD intensity at 222 nm. CD spectra in far UV region for P5 and P5A5 measured at different temperatures (left), and the molar ellipticity at 222 nm plotted as a function of temperature (right). **b,** Disulfide-linked oligomerization of P5 and its mutants in the presence of various molar ratios of reduced and oxidized glutathione (GSH/GSSG). The lower panel shows quantification of the oligomer fractions (including a dimer) shown in the upper panels (n = 3; mean ± SD). The green region indicates the redox condition corresponding to that in the ER.

**Table 4.** Thermodynamic parameters for heat denaturation of P5 and P5A5.

| | *T<sub>m</sub>*<br>(°C) | $\Delta H_{\text{unfold}}$<br>(kJ/mol) |
| --- | --- | --- |
| **P5** | 55.3±0.4 | 585.7±32.3 |
| **P5A5** | 49.1±0.9 | 230.3±46.2 |

To investigate whether structural instability causes other anomalies in P5, we examined the possible generation of aberrant disulfide-linked P5 oligomers under a wide range of redox conditions. Consequently, we observed that disulfide-linked oligomers were significantly generated for P5A5, but not for P5, under ER-like redox conditions of [GSH]^2^/[GSSG] = 0.5–3 mM (Fig. 5b, and Extended Data Fig. 9)(Hwang, Sinskey et al., 1992). To further explore whether the **a^0^** or **a** domain active site of P5A5 is involved in the disulfide-linked oligomers, the redox-active site cysteines in either domain were replaced by alanine to prepare P5A5 AXXA-CXXC (AC) and CXXC-AXXA (AC) mutants. Disulfide-linked oligomers were observed for P5A5-CA similarly to P5A5, but this was not the case for P5A5-AC or the active-site mutants of P5 (P5-CA and P5-AC) (Fig. 5b, and Extended Data Fig. 9), indicating that the disulfide-linked oligomers were formed via domain **a^0^** of P5A5. Thus, loss of the Leu-Val adhesive motif causes conformational destabilization within the P5 **a^0^** domain.

### Dimerization via the Leu-Val adhesive motif stabilizes a Trx-like fold of domain a^0^

The above results demonstrated that the Leu-Val adhesive motif in domain **a^0^** is critical for stabilizing the overall structure of P5, and for suppressing the formation of artefactual disulfide-linked oligomers. To further explore structural alterations caused by the loss of the motif, we utilized nuclear magnetic resonance spectroscopy (NMR). The ^1^H-^15^N hetero-nuclear single quantum coherence (HSQC) spectrum of the uniformly ^15^N-labaled **a^0^** domain showed dispersed resonances, indicating that **a^0^** adopts a folded structure in solution (Fig. 6a). The HSQC spectrum of the **a^0^**A5 mutant also showed the dispersed resonances whose chemical shift roughly matched to those of **a^0^**. Thus, the overall structure of **a^0^** has been almost maintained in the mutant (Fig. 6a). However, the spectra showed significant chemical shift changes for several resonances upon the introduction of A5 mutations (Fig. 6a and 6b). Chemical shift perturbations due to the loss of the Leu-Val adhesive motif were observed not only for residues of α-helix 4 located at the dimer interface, but also for residues of α-helix 2 in close contact with α-helix 4, and for residues of the β-sheet buried inside the molecule (Fig. 6b).

**Fig. 6.**
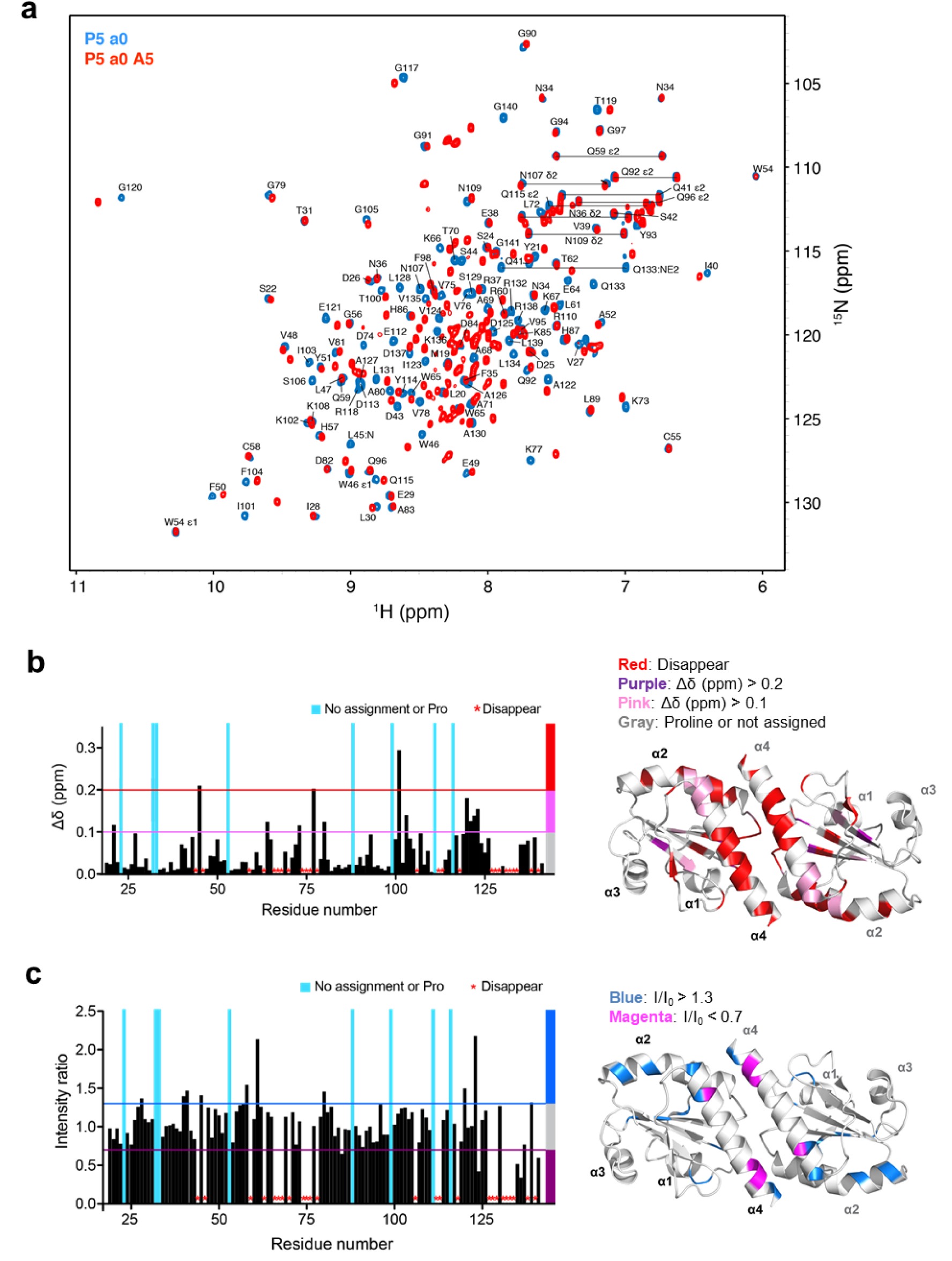
Loss of the Leu-Val adhesive motif causes local unfolding of the P5 a^0^ domain. **a,** Overlay of 2D ^1^H-^15^N HSQC spectra of **a^0^** and **a^0^** with the mutations of A5 (**a^0^**A5). **b, (**Left) Residue-specific chemical shift difference between **a^0^** and **a^0^**A5. (Right) Mapping of the chemical shift difference between **a^0^** and **a^0^**A5 onto the ribbon representation of the crystal structure of **a^0^**. Red, purple, pink, and gray indicate residues of which the signals disappear or display significant chemical shift deviations upon the mutations of A5; signals disappear, Δδ (ppm) >0.2, Δδ (ppm) >0.1, and Pro/no assignment, respectively. **c, (**Left) Residue-specific intensity changes of signals in the ^1^H-^15^N HSQC spectra of **a^0^** and **a^0^**A5. (Right) Blue and magenta indicate residues of which the signals display significant intensity changes upon the mutations of A5; I/I0 >1.3 or I/I0 <0.7, respectively.

In the spectrum of **a^0^**A5, several resonances disappeared, whereas several additional resonances appeared around the center of the spectrum (^1^H 7.5 ∼ 8.5 ppm). Given the fact that the resonances located around the center of the spectrum are indications of disordered regions of the protein, the spectrum suggests that the regions including α-helix 2 and 4 are destabilized and partially disordered in **a^0^**A5. Mapping of the intensity ratio supported the partial disorder of **a^0^**A5: In addition to the chemical shift changes, intensity changes were also observed especially for the resonances corresponding to residues of α-helix 2 (Fig. 6c). The increased intensity of the resonances for these specific residues can be explained by the increased dynamics or disorder of α- helix 2 upon the disruption of the dimer interface and the loosely packed fold of **a^0^**A5, in agreement with the lower thermal stability of P5A5 than that of P5 (Fig. 5a).

## Discussion

In the present study, we discovered a novel adhesive motif in the N-terminal Trx-like domain of P5, and found that this motif stabilizes the dimeric structure of P5 and ensures its disulfide reductase activity and Ca^2+^-dependent chaperone function. Conventional Leu-zipper motifs, which are frequently found at the dimer interface of DNA binding proteins including transcription factors(Landschulz et al., 1988), also stabilize the dimeric structures, but have different function such as grabbing double-stranded DNA near the ends of two α-helices included in this motif to regulate gene expression.

Detailed structure comparison between these two dimerization motifs revealed that at the dimer interface of P5, two regions separately located on the primary structure (region 1 and region 2) contribute to dimerization, whereas a single broader region is involved in the dimerization of Leu-zipper motifs (Fig 7a and b). Based on the *in silico* calculation with a program Protein Interfaces, Surfaces and Assemblies (PISA)(Krissinel & Henrick, 2007), the major adhesive force in the P5 dimer derives from region 2 corresponding to α-helix 4, although the contribution of region 1 corresponding to α-helix 2 is not negligible (Fig. 7a). Consistently, significant chemical shift changes were observed for residues located at α-helix 4 and α-helix 2 in the HSQC spectrum of **a^0^**A5 (Fig. 6). The *in silico* calculation also demonstrated that binding free energy (Δ*G*) per buried surface area for the P5 adhesive motif is ca. -10 ×10^-3^ (kcal/mol/Å^2^), whereas those for typical Leu-zipper motifs range from -15 to -20 ×10^-3^ (kcal/mol/Å^2^) (Fig. 7c). The Δ*G*/residue is calculated to be ca. -0.4 for the P5 adhesive motif and ca. -0.8 (kcal/mol) for the Leu zipper motifs, respectively. The two-dimensional plots of Δ*G*/area versus Δ*G*/residue clearly shows that the P5 adhesive motif provides significantly lower binding free energy or weaker adhesive force than Leu-zipper motifs (Fig. 7d). Thus, these two sorts of motifs are interpreted to work with different modes of interaction and different mechanical stability while commonly serving to promote protein dimerization.

**Fig. 7.**
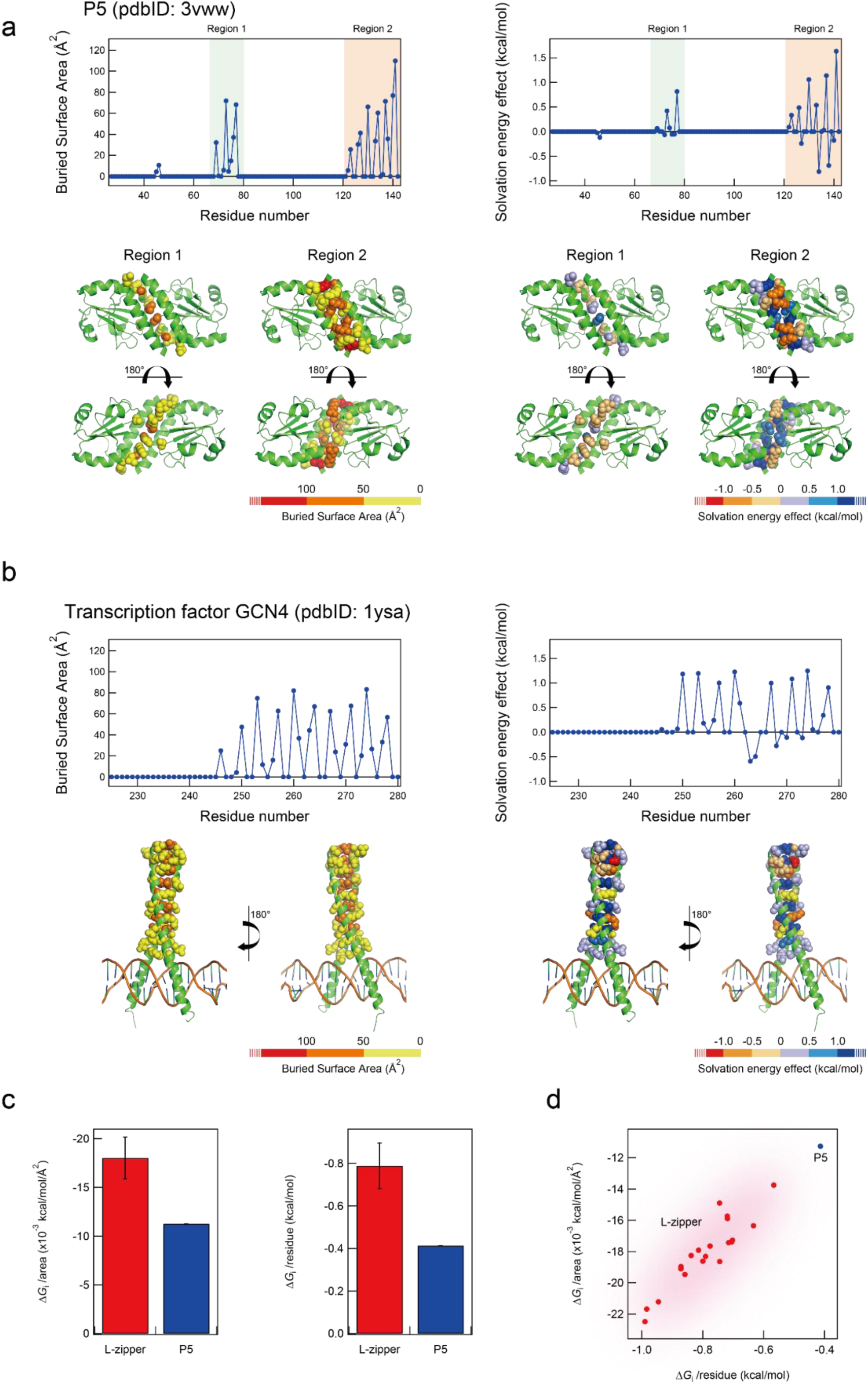
*In silico* analysis for the binding free energies of the P5 Leu-Val adhesive motif and of typical Leu-zipper motifs found in DNA binding proteins. **a**, *In silico* analysis was carried out using a program Protein Interfaces, Surfaces and Assemblies (PISA). Left and right panels represent the buried surface area and solvation energy effect of each residue for the P5 adhesive motif. **b,** Results of the same *in silico* analysis as described in **a** for the Leu-zipper motif of transcription factor GCN4 (PDB code: 1ysa). **c**, Comparison of the binding free energy (Δ*G*) per buried surface area and per residue between Leu-zipper motifs and the adhesive motif in P5. Note that the values for the Leu-zipper are calculated based on 20 kinds of typical Leu-zipper motifs seen in DNA binding proteins of know structure. **d,** Two-dimensional plots of Δ*G*/area versus Δ*G*/residue.

Regarding the structure-function relationship of P5, the protein comprises two redox-active and one redox-inactive Trx-like domains to play important roles in oxidative and reductive processes in the ER(Eletto et al., 2016, Kaiser et al., 2007, Sato et al., 2013). The present SAXS analysis revealed that P5 forms a homodimer of which the six Trx-like domains are separately arranged in multiple possible ways and that both the redox-active sites in domains **a^0^** and **a** are exposed to the solvent. The presence of multiple solvent-exposed redox-active sites in the mobile Trx-like domains may be advantageous for P5 to rapidly and promiscuously introduce disulfide bonds into unfolded substrates due to the easier access to any pair of cysteines on substrates. In contrast to those of P5, the redox-active sites of PDI are located such that they face each other across a central substrate-binding cleft in the overall U-like shape. Such well organized locations of the redox-active sites in PDI seem suitable for the enzyme to act on partially folded substrates with compact structures by binding them via the central cleft and catalyzing thiol-disulfide exchanges via the two redox-active sites facing each other.

While some physiological significance of P5 dimerization has been clarified in this work, the monomeric P5 mutant can introduce disulfide bonds into substrates as effectively as P5 (Fig. 2c). Regarding the dimerization of PDI family members, we recently demonstrated that, in the presence of unfolded substrates, PDI assembles to form non-covalent dimers and accelerates oxidative folding of substrates inside the central cavity created by the PDI dimers (Okumura, Noi et al., 2020, Okumura et al., 2019). Unlike the substrate-induced PDI dimers, P5 dimer is likely to form constitutively even without substrates and does not appear to create a cavity for binding substrates tightly. The dimeric conformation with variable Trx-like domain arrangements will provide high adaptability to substrates with different sizes and shapes. Monomerization-induced partial unfolding of P5 (Fig. 6) may lead to looser binding or recognition to substrates, especially those with folded structures like the SS-linked IRE1 LD oligomer and insulin, possibly explaining the compromised disulfide reduction activity of P5A5 (Fig. 3).

For the maintenance of ER homeostasis, various molecular chaperones work cooperatively, and many of them, including PDIs, consist of multiple domains and/or form oligomers. To acquire sophisticated function without causing folding problems, fusion of multiple domains and assembly of multiple protomers could be a simple and effective strategy(Han, Batey et al., 2007). In this regard, it is noteworthy that Trx-like domains commonly shared by many proteins including even redox-irrelevant proteins have evolved an adhesive motif essential for dimerization and structural stabilization in P5, a ubiquitously expressed PDI family enzyme involved in the ER quality control. There could be yet-unidentified adhesive motifs that promote protein dimerization or higher-order oligomerization for fulfilling important physiological function. Extensive structural and bioinformatics studies will further reveal unique modes of interaction that ensure structural stability and protein multimerization and thereby support or upgrade protein function.

## Acknowledgments

Synchrotron radiation experiments were performed on beamline BL45XU in SPring-8 with the approval of RIKEN (Proposal No. 2014A1345, 2017B1176, 2018A1311, 2018B1457). This work was partly supported by Nanotechnology Platform Program <MOLECULE and Material Synthesis> (JPMXP09S20MS0001) of the Ministry of Education, Culture, Sports, Science and Technology (MEXT), Japan. We are grateful to N. Fukamachi (Tohoku University) for experimental assistance. We are grateful to Dr. Y. Lin from KBSI for ITC analysis. We thank Dr. Kozo Tanaka (Tohoku University) for providing HeLa Kyoto cells.

## Financial support

This research was funded by JSPS KAKENHI Grant Number JP17H06521 (to S.K.), JP19K16092 (to S.K.), JP19J00893 (to M.M.), and JP20K15969 (to M.M.). We acknowledge, with thanks, funding from a Grant-in-Aid for Scientific Research on Innovative Areas from MEXT (19H04799 and 20H04688 to M.O.), the Takeda Science Foundation (to K.I. and M.O.), the Mochida Memorial Foundation for Medical and Pharmaceutical Research (to M.O.), the Japan Foundation of Applied Enzymology (to M.O.), the Building of Consortia for the Development of Human Resources in Science and Technology (to M.O.), a Grant-in-Aid for Scientific Research (C) to M.O. (19K06520) and Scientific Research (A) to KI (18H03978), the Promotion of Joint International Research (Fostering Joint International Research (B)) to T.S., M.O., and S. K. (20345793), Ensemble Grants for Early Career Researchers 2020 to M.M. and M.O., the National Research Foundation of Korea (NRF) grants funded by the Korean government [NRF-2018K1A3A1A39088040 and NRF-2019R1A2C1004954 (Y.-H.L.)], the National Research Council of Science & Technology (NST) grant funded by the Korea government (MSIP) (CAP-17-05-KIGAM) (Y.-H.L.), and the Korea Basic Science Institute grant (C070410) (Y.-H.L.).

## Author contributions

M.O. designed and performed parts of experiments, including SAXS and oxidative protein folding experiments. S.K. performed SAXS and chaperone activity experiments. M.M. prepared purified IRE1 LD, and performed reduction assays and P5 overexpression experiments in cultured cells. M.K., D.I, and Y.-H. L. performed ITC and CD experiments. T.S. and H.K. performed NMR experiments. C.H. and M.W. prepared P5 and P5 mutant proteins. Y.A. assisted with cell culture experiments. S.A. analyzed the SAXS data. K.I. supervised the study. K.I. and M.O. wrote the manuscript. M.O. prepared figures. All authors discussed the results and approved the manuscript.

---

**Extended Data Fig. 1.**
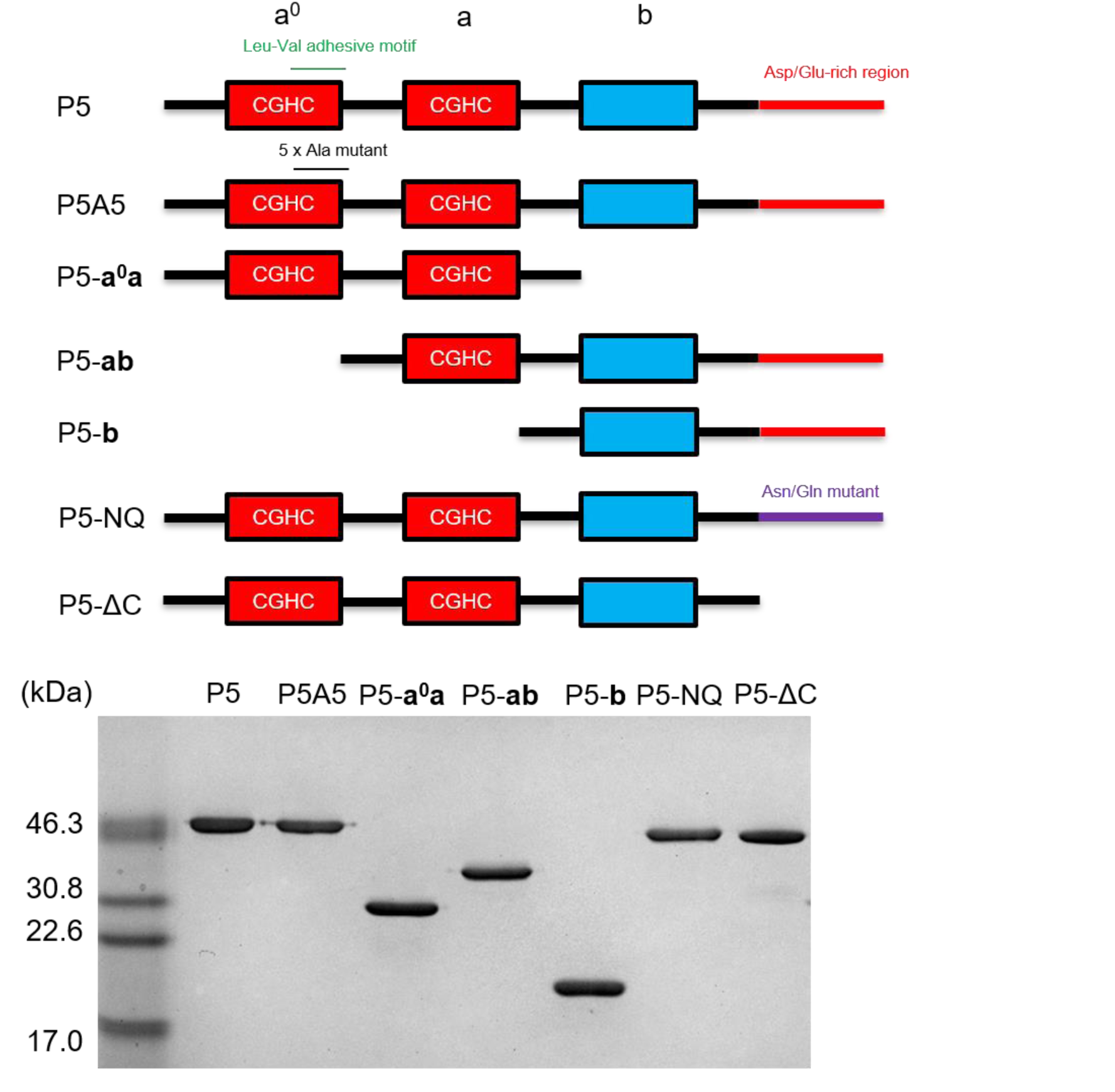
Non-reducing SDS-PAGE analysis of P5 and its mutants constructed in this study. (upper) Schematic structures of P5 and its mutants constructed in this study. Trx-like domains are indicated by red and blue boxes, respectively. Active site sequences in the redox-active domains are indicated in red boxes. (lower) Non-reducing SDS-PAGE analysis verified that all constructs ran as single bands and were fully purified.

**Extended Data Fig. 2.**
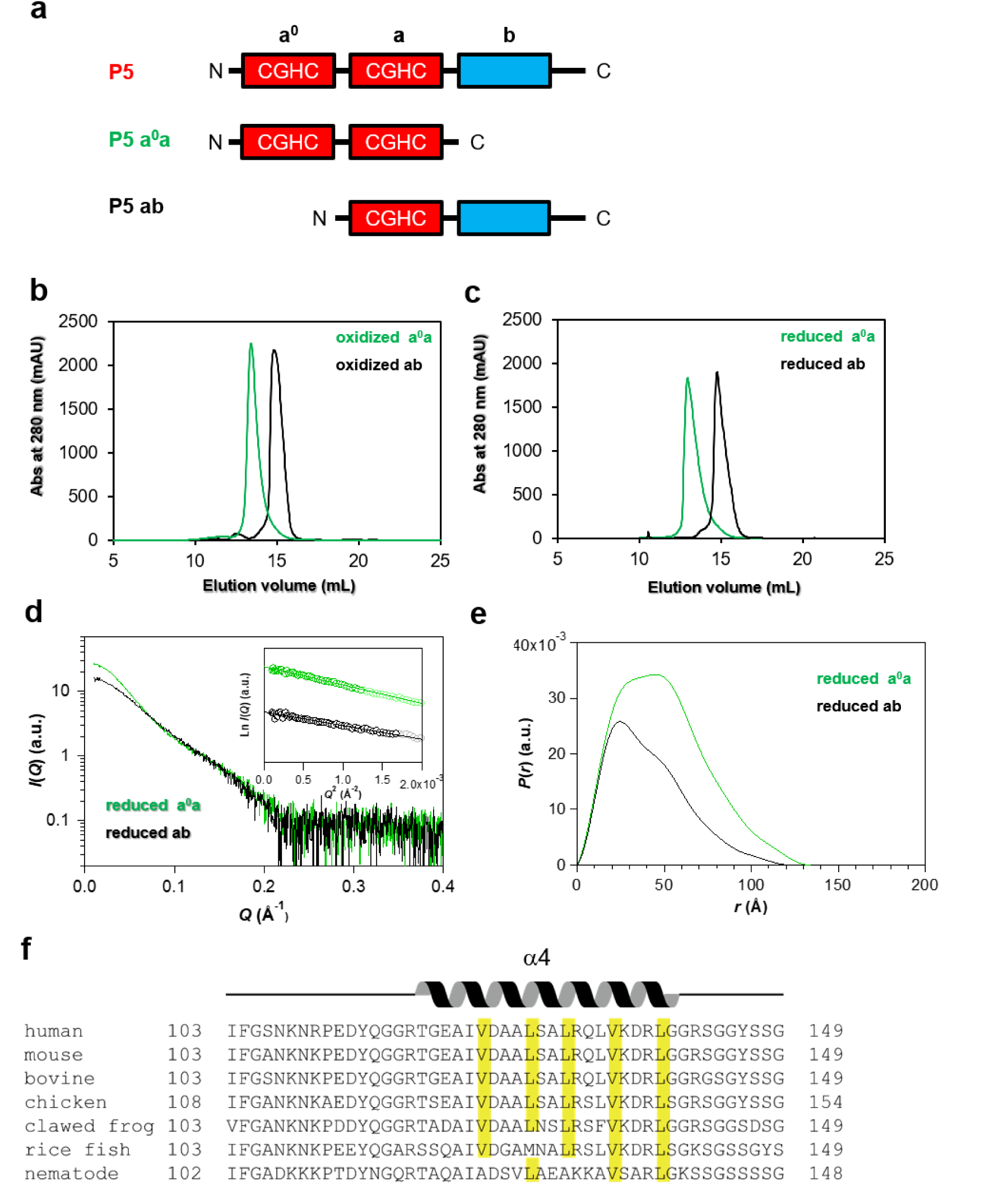
P5 dimerizes via the a^0^ domain. **a,** Domain organization of P5, P5 **a^0^a** and P5 **ab**. Redox-active and redox-inactive Trx-like domains are indicated by red and blue boxes, respectively. Active site sequences in the redox-active domains are indicated in red boxes. **b** and **c,** Size-exclusion chromatography analysis of oxidized (**b**) and reduced (**c**) **a^0^a** or **ab**. **d,** SAXS profiles of reduced **a^0^a** (green) and **ab** (black). The inset shows Guinier plots using the *Q* range (highlighted data points in the inset) shown in Table 1. **e,** Pair distribution function *P*(*r*) of reduced **a^0^a** (green) and **ab** (black). **f,** Alignment of the amino acid sequence around α-helix4 of P5 from different species. Highly conserved leucine and valine residues contained in the adhesive motif are highlighted in yellow.

**Extended Data Fig. 3.**
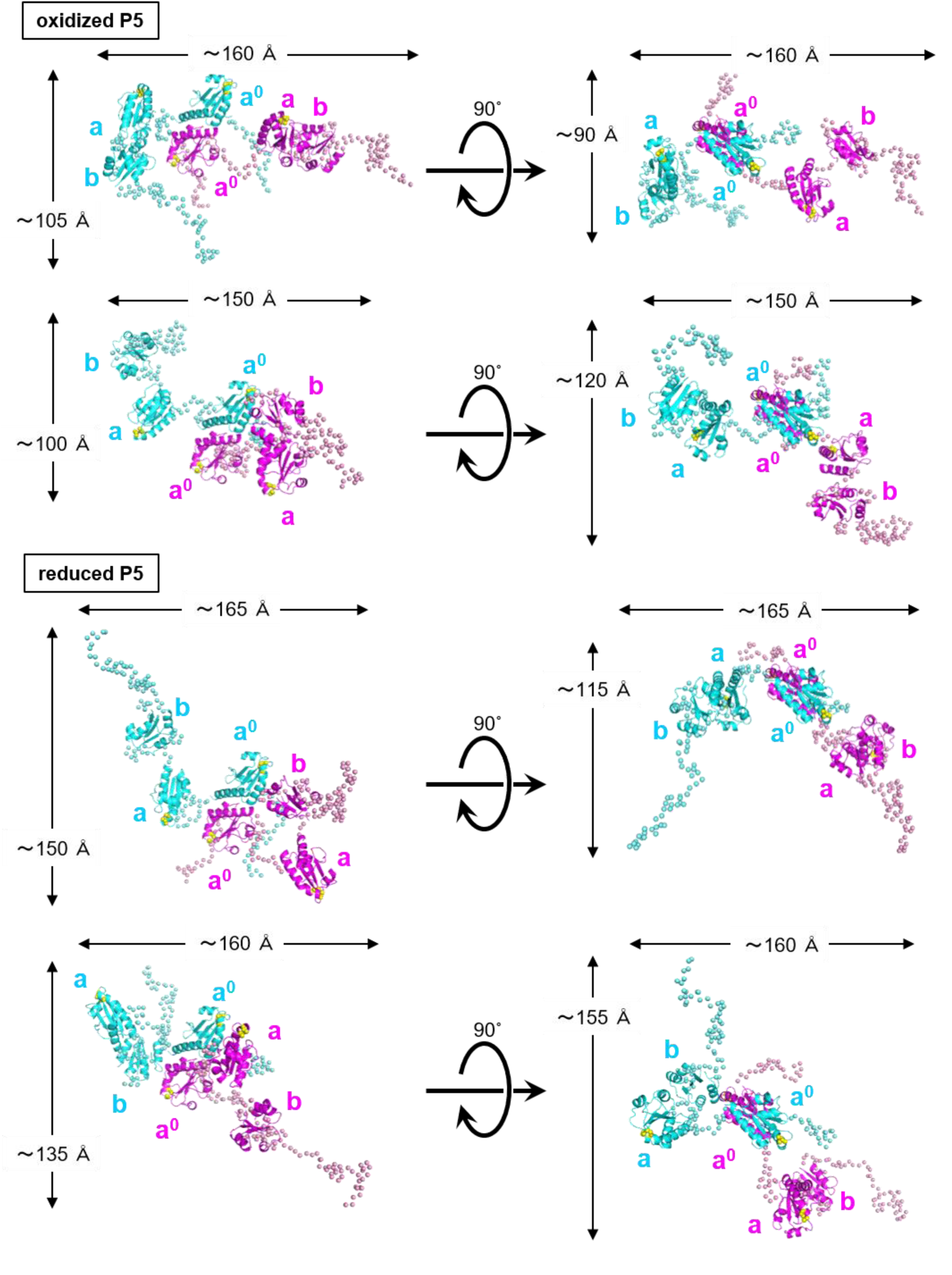
Feasible SAXS structure models of oxidized (upper) and reduced (lower) forms of P5. Multiple rigid-body refinement models of oxidized (upper) and reduced (lower) forms of P5 generated using CORAL are represented by a ribbon diagram. Redox active sites in domains **a^0^** and **a** are shown by yellow spheres. (see Table 2 for more details)

**Extended Data Fig. 4.**
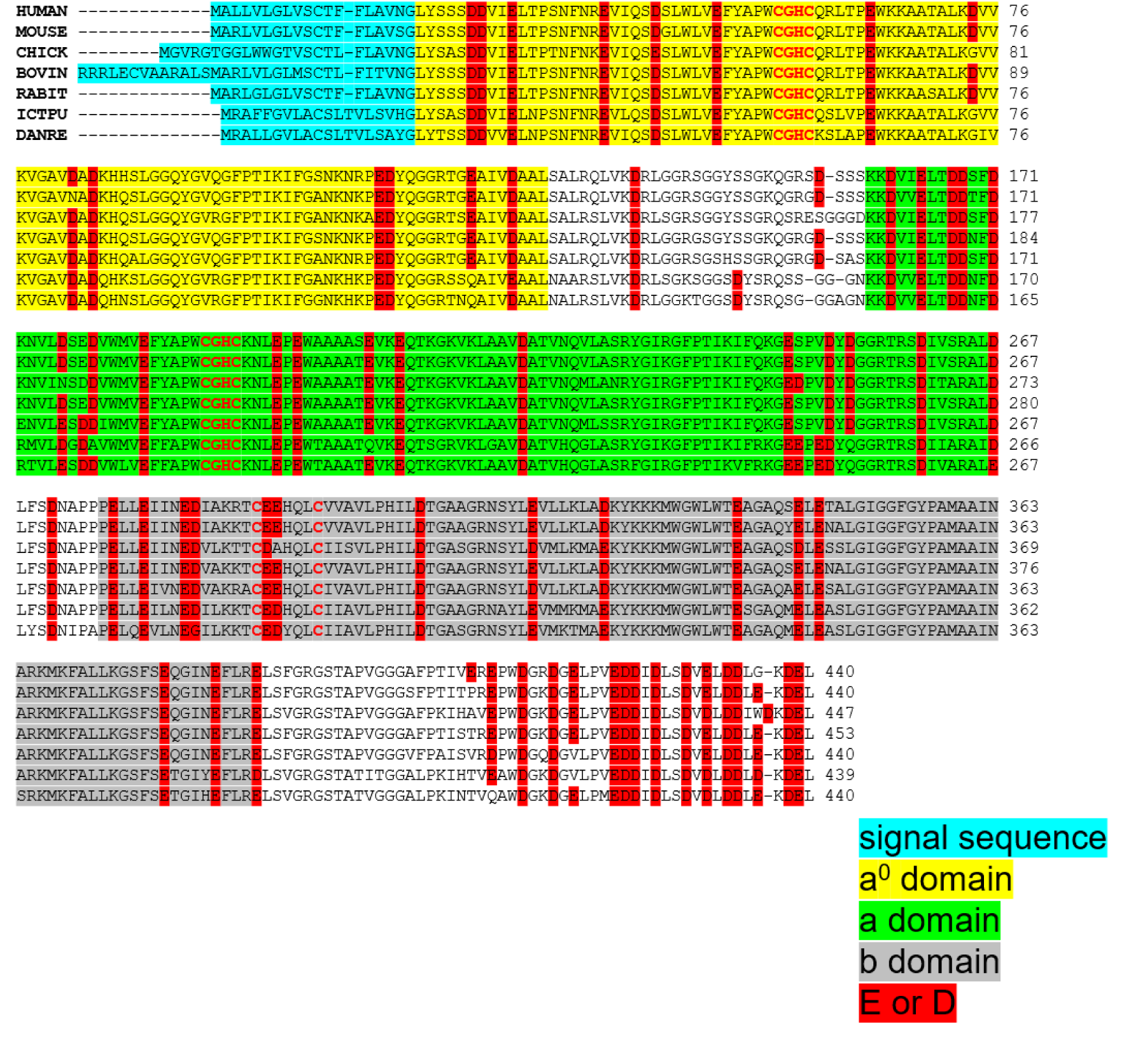
Alignment of the amino acid sequences of P5 from different species. Redox-active sites in domains **a^0^** and **a** are denoted by red letters. Highly conserved aspartate and glutamate are boxed in red.

**Extended Data Fig. 5.**
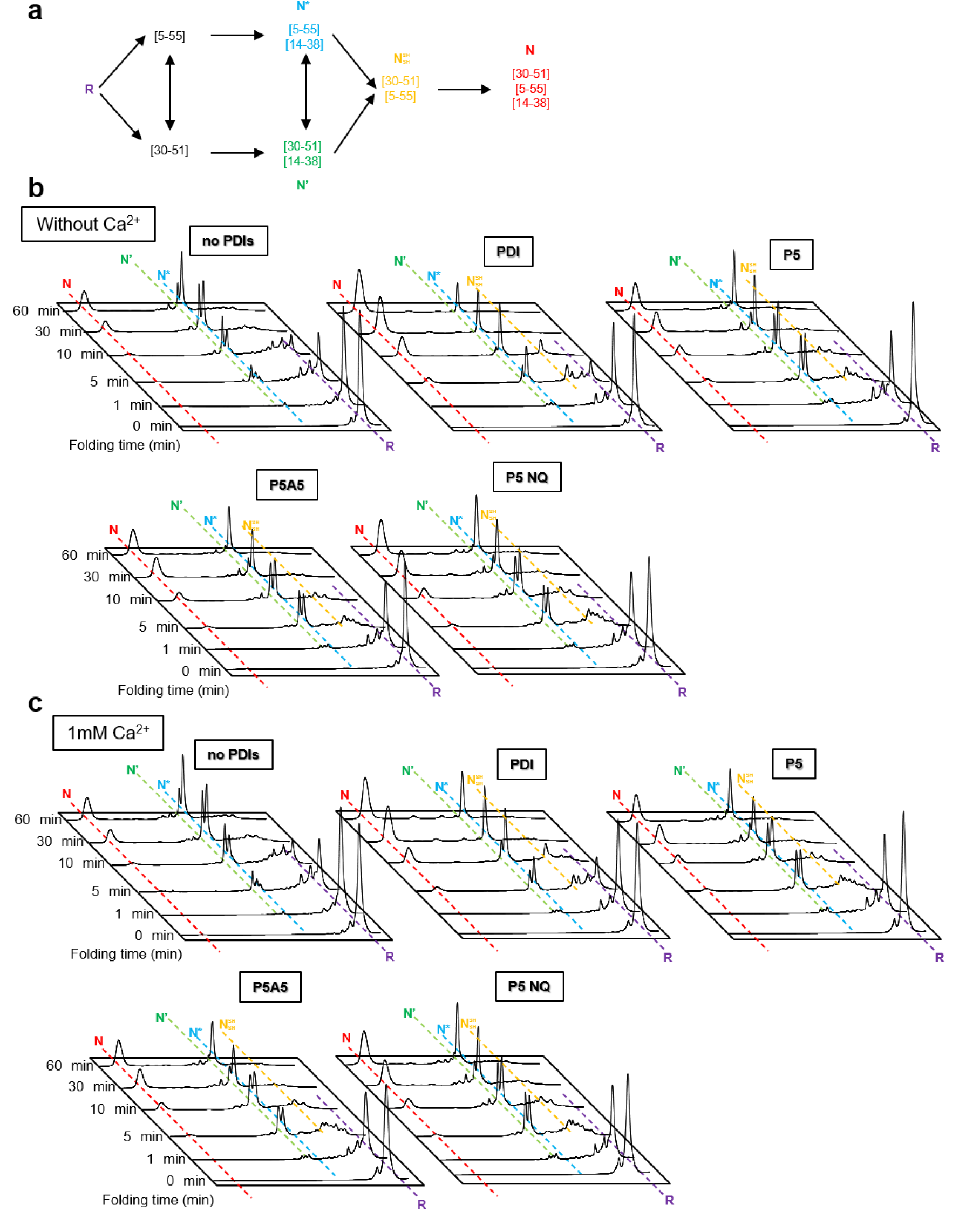
Oxidative folding of BPTI catalyzed by PDI, P5, and P5 mutants. **a,** Schematic representation of disulfide formation pathways during BPTI folding(Weissman & Kim, 1991). R, 1SS, 2SS, and N indicate the reduced, one disulfide-bonded, two disulfide-bonded, and native species, respectively. N represents native BPTI with [5–55; 30–51; 14–38]; N^SH^, N*, and N’ represent BPTI folding intermediates with two disulfide bonds, [30–51; 5–55], [5–55; 14–38], and [30–51; 14–38], respectively. **b,** HPLC profiles showing the time course of oxidative folding of BPTI catalyzed by no enzyme (control), PDI, P5, and P5 mutants in the absence of Ca^2+^. At the indicated time points, reaction mixtures were quenched with HCl, and analyzed by HPLC and MALDI-TOF/MS. The same trend was observed in all three independent experiments. **c,** HPLC profiles showing the time course of oxidative folding of BPTI catalyzed by no enzyme (control), PDI, P5, and P5 mutants in the presence of Ca^2+^. The same trend was observed in all three independent experiments.

**Extended Data Fig. 6.**
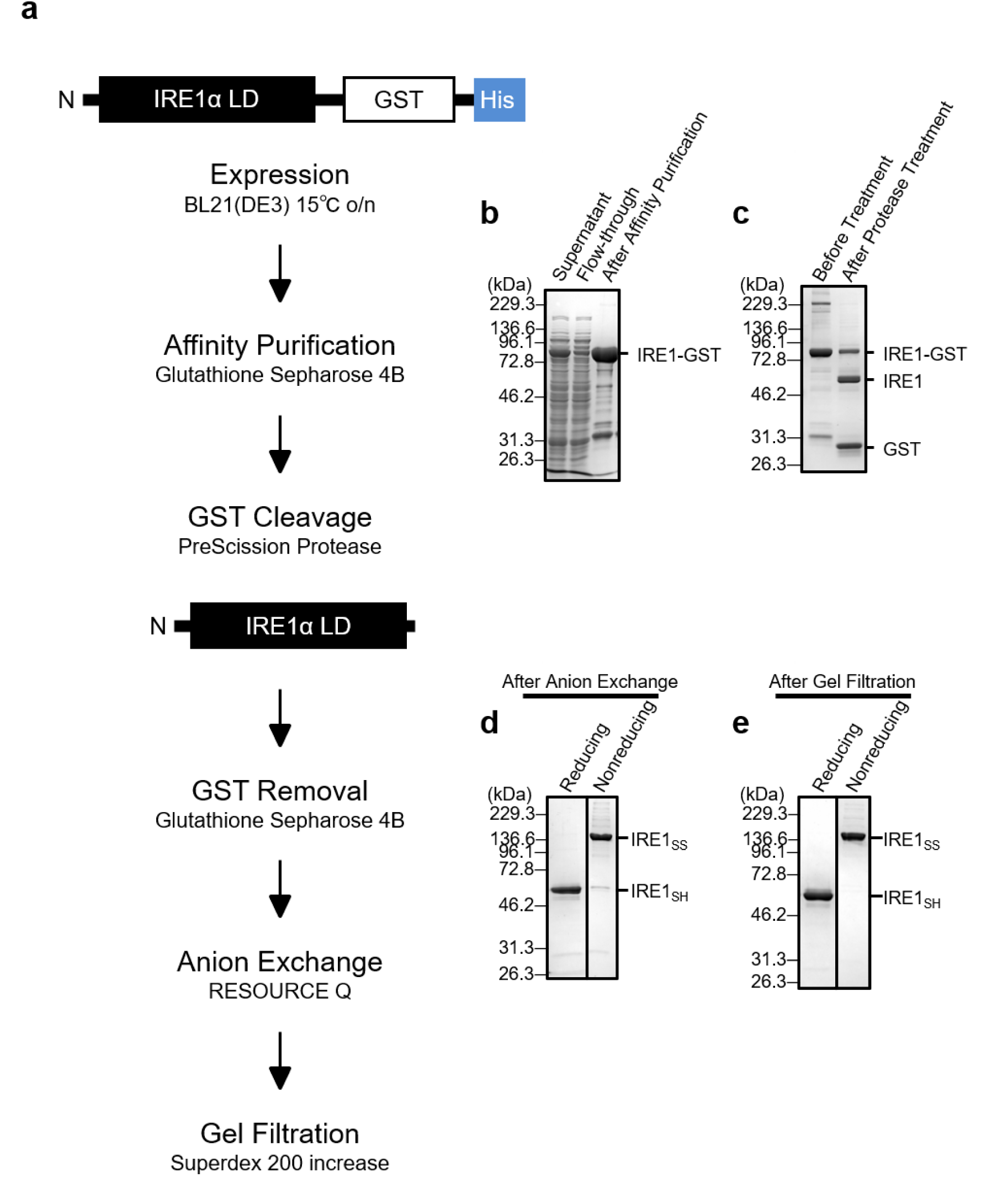
Expression and purification of the IRE1α luminal domain. **a,** Schematic representation of the expression and purification of the IRE1α luminal domain. IRE1α luminal domain (LD) without a signal peptide was fused with glutathione S-transferase (GST) and a His6-tag at the C-terminus (IRE1α LD-GST), overexpressed in *E. coli* strain BL21 (DE3), and purified as described in the Materials and Methods. **b,** SDS-PAGE analysis of IRE1α LD-GST before and after GST-affinity purification. IRE1-GST, fusion protein of IRE1α LD and GST. **c**, SDS-PAGE analysis of IRE1α LD-GST before and after protease treatment. IRE1 and GST represent GST-cleaved IRE1α LD and the cleaved-off GST, respectively. **d,** SDS-PAGE analysis of IRE1α LD after anion exchange chromatography. IRE1SS and IRE1SH represent disulfide (SS)-linked IRE1 LD and reduced IRE1α, respectively. **e,** SDS-PAGE analysis of IRE1α LD after size-exclusion chromatography.

**Extended Data Fig. 7.**
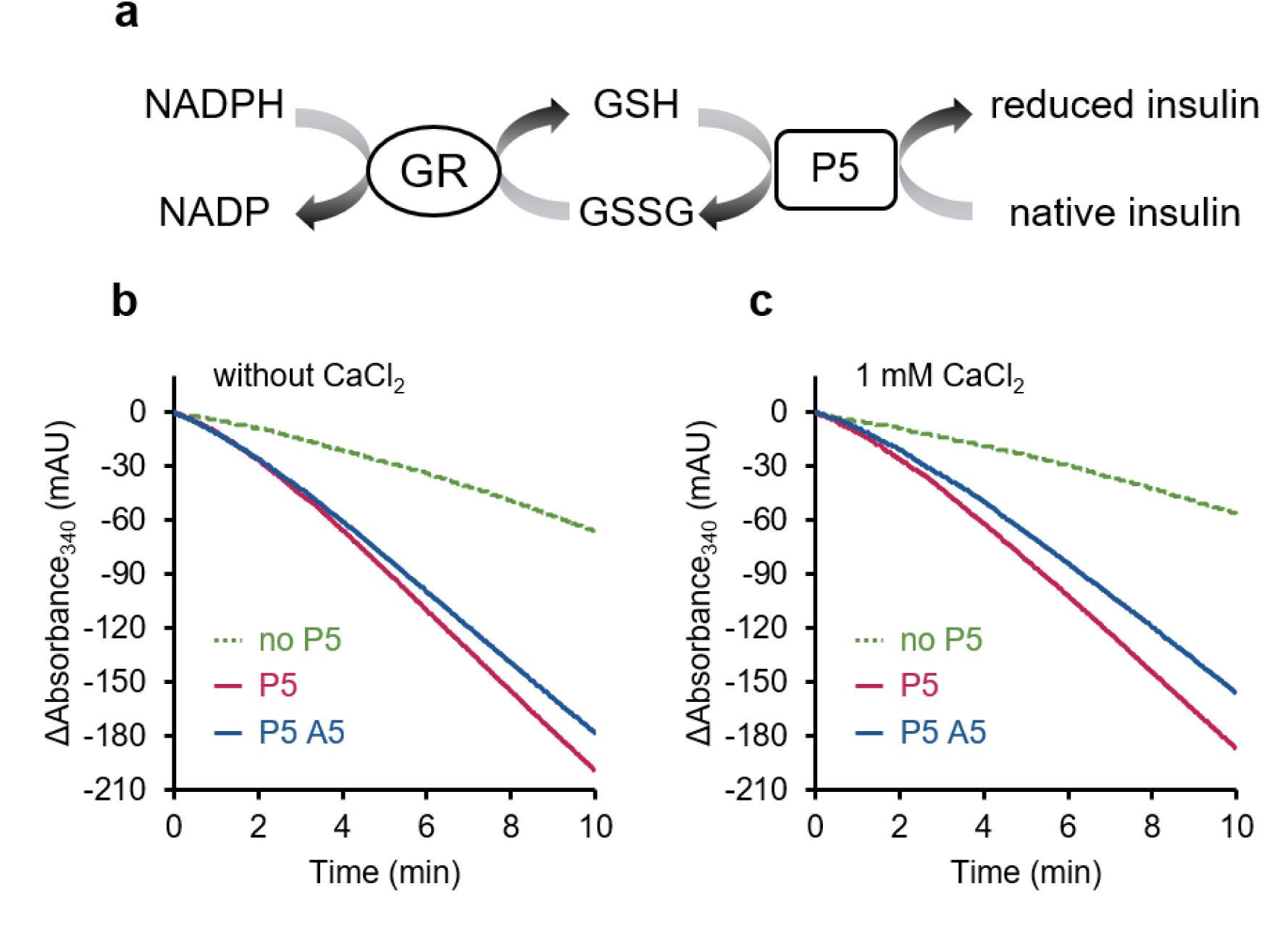
Insulin reduction assay of P5 and P5A5 with and without _Ca_2+. **a,** Schematic representation of P5-mediated insulin reduction coupled with the glutathione reduction by glutathione reductase (GR) and NADPH. **b,** NADPH consumption with or without 1 µM P5 or P5A5 in the presence of 30 μM insulin, 10 mM GSH, 200 μM NADPH, and 2 U/mL glutathione reductase. NADPH consumption was monitored as the change in absorbance at 340 nm (mAU, ×10^-3^ arbitrary units). **c,** Results of the same experiments as described in **b** but with 1 mM CaCl2.

**Extended Data Fig. 8.**
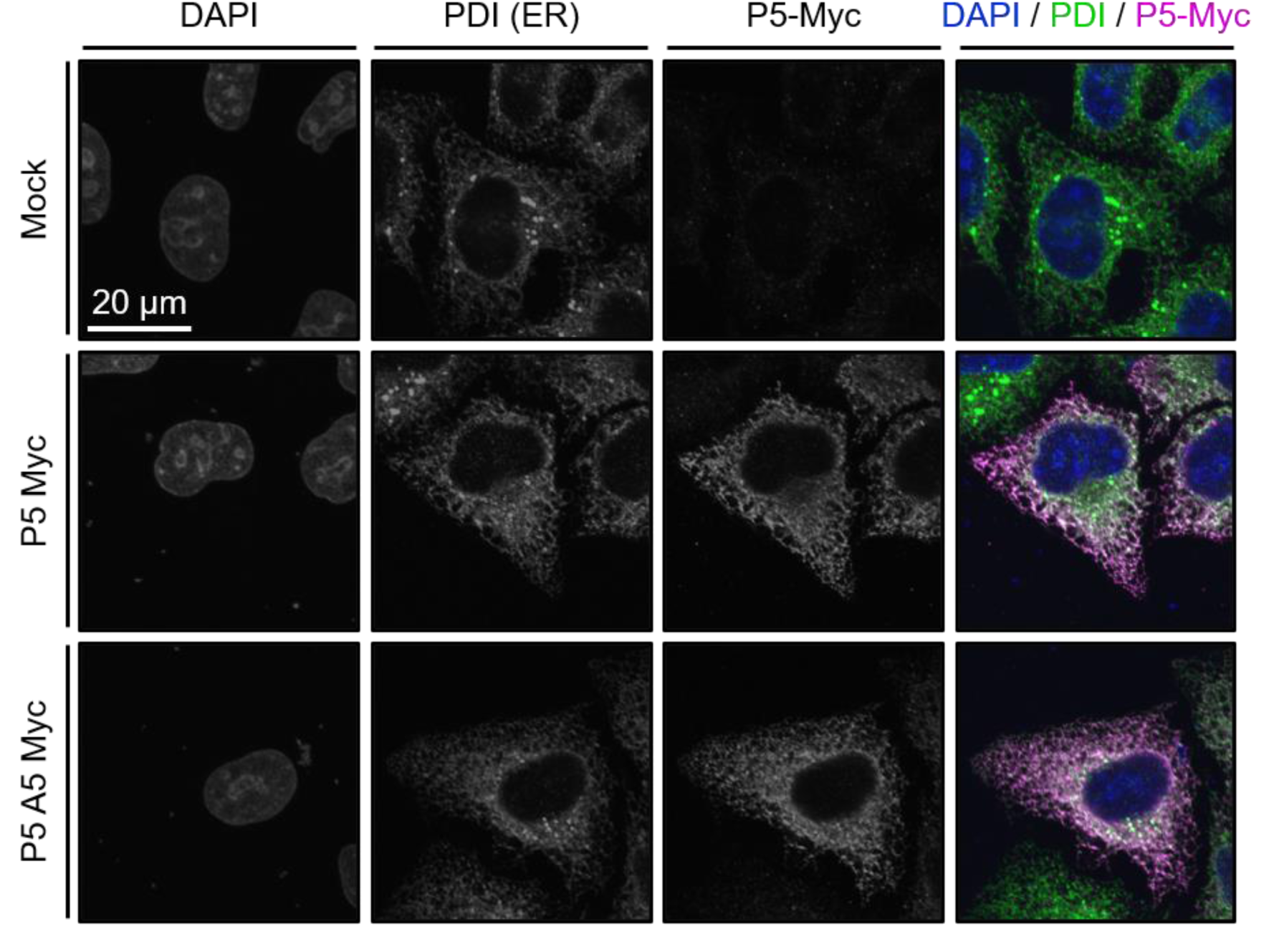
Subcellular localization of exogenously expressed P5. Confocal immunofluorescence images showing the subcellular localization of Myc-tagged P5 (magenta) in HeLa Kyoto cells. Cells were co-stained with an antibody recognizing PDI (green) and with DAPI (blue) to highlight the ER and nucleus, respectively. Note that P5 and P5A5 are localized in the ER.

**Extended Data Fig. 9.**
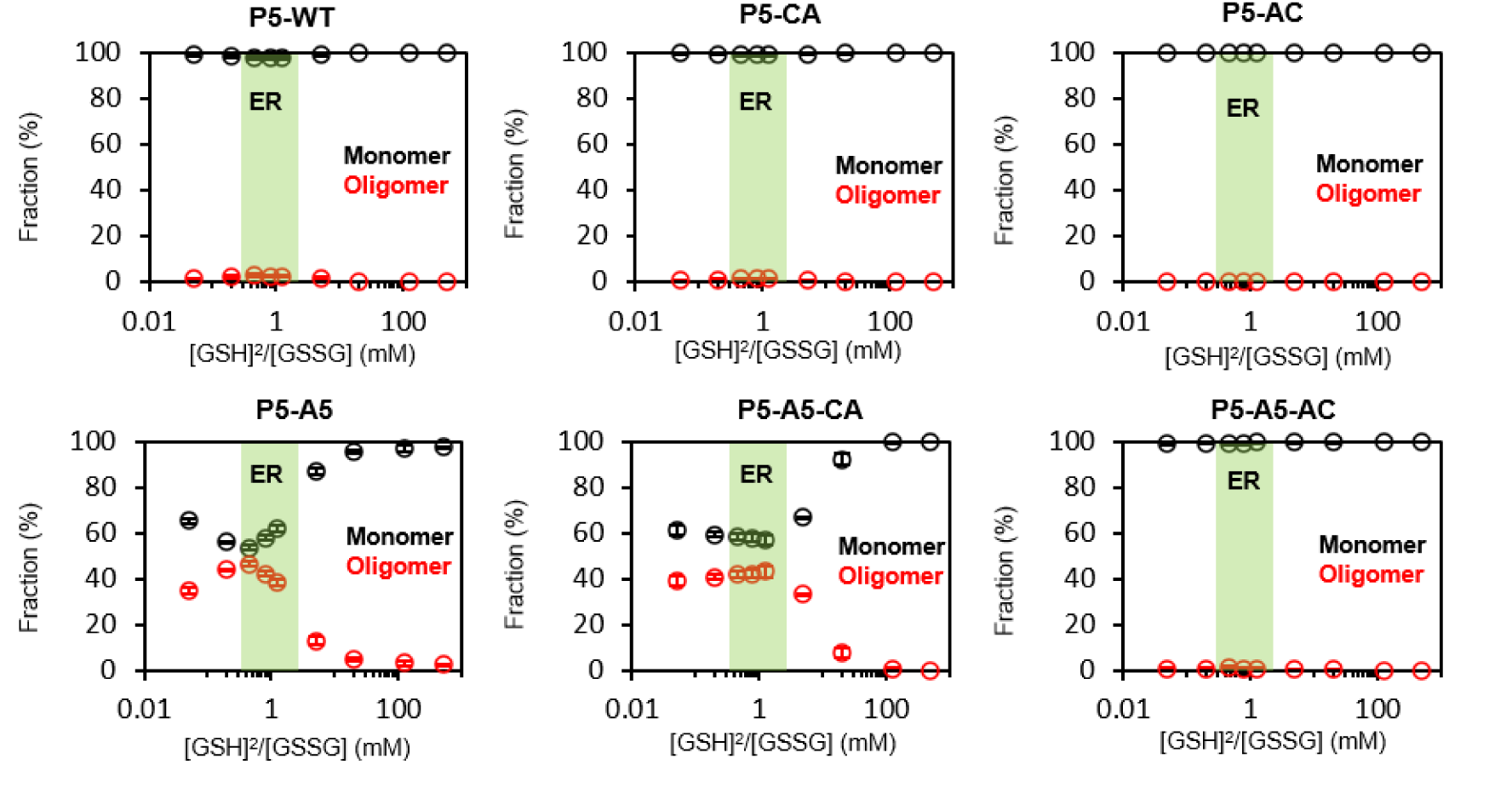
Monomer and disulfide-linked oligomer species of P5 and its mutants generated under various redox conditions. Fractions of monomer and disulfide-linked oligomer (including a dimer) species of P5 and its mutants generated under various redox conditions were quantified and plotted as a function of different ratios of reduced and oxidized glutathione. The green region indicates the redox condition corresponding to that of the ER.

**Supplementary Table 1.** Plasmid DNA.

| No. | plasmid name | vector | insertion site | tag | insertion (protein) | mutation position |
| --- | --- | --- | --- | --- | --- | --- |
| 1 | **P5-a0ab (WT: CxxC, CxxC)** | pET15b (Novagen) | Nde I-BamH I | His<sub>6</sub> at N-term | 20-440 a.a.<br>(1-19: signal sequence) |  |
| 2 | **P5-a0ab (AxxA, CxxC)** | pET15b (Novagen) | Nde I-BamH I | His<sub>6</sub> at N-term | 20-440 a.a.<br>(1-19: signal sequence) | Cys55Ala/Cys58Ala |
| 3 | **P5-a0ab (CxxC, AxxA)** | pET15b (Novagen) | Nde I-BamH I | His<sub>6</sub> at N-term | 20-440 a.a.<br>(1-19: signal sequence) | Cys190Ala/Cys193Ala |
| 4 | **P5-a0a** | pET15b (Novagen) | Nde I-stop codon | His<sub>6</sub> at N-term | 20-290 a.a.<br>(1-19: signal sequence) | stop codon at 291 |
| 5 | **P5-a0** | pET15b (Novagen) | Nde I-stop codon | His<sub>6</sub> at N-term | 20-141 a.a.<br>(1-19: signal sequence) | stop codon at 142 and 143 |
| 6 | **P5-ab** | pET15b (Novagen) | Nde I-BamH I | His<sub>6</sub> at N-term | 141-440 a.a. | deletion of 20-140 |
| 7 | **P5-b** | pET15b (Novagen) | Nde I-BamH I | His<sub>6</sub> at N-term | 273-440 a.a. | deletion of 20-272 |
| 8 | **P5-Δc** | pET15b (Novagen) | Nde I-BamH I | His<sub>6</sub> at N-term | 20-421 a.a.<br>(1-19: signal sequence) | deletion of 422-440 |
| 9 | **P5-NQ** | pET15b (Novagen) | Nde I-BamH I | His<sub>6</sub> at N-term | 20-440 a.a.<br>(1-19: signal sequence) | E422Q, D423N, D424N,<br>D426N, D429N, E431Q,<br>D433N, D434N |
| 10 | **P5-V124A** | pET15b (Novagen) | Nde I-BamH I | His<sub>6</sub> at N-term | 20-440 a.a.<br>(1-19: signal sequence) | V124A |
| 11 | **P5-V124A/L128A** | pET15b (Novagen) | Nde I-BamH I | His<sub>6</sub> at N-term | 20-440 a.a.<br>(1-19: signal sequence) | L128A |
| 12 | **P5-V124A/L128A/L139A** | pET15b (Novagen) | Nde I-BamH I | His<sub>6</sub> at N-term | 20-440 a.a.<br>(1-19: signal sequence) | L139A |
| 13 | **P5-V124A/L128A/V135A/L139A** | pET15b (Novagen) | Nde I-BamH I | His<sub>6</sub> at N-term | 20-440 a.a.<br>(1-19: signal sequence) | V135A |
| 14 | **P5-A5**<br>**(V124A/L128A/L131A/V135A/L139A)** | pET15b (Novagen) | Nde I-BamH I | His<sub>6</sub> at N-term | 20-440 a.a.<br>(1-19: signal sequence) | L131A |
| 15 | **P5-a0 (A5)** | pET15b (Novagen) | Nde I-stop codon | His<sub>6</sub> at N-term | 20-141 a.a.<br>(1-19: signal sequence) | stop codon at 142 and 143 |
| 16 | **P5-A5 (AxxA, CxxC)** | pET15b (Novagen) | Nde I-BamH I | His<sub>6</sub> at N-term | 20-440 a.a.<br>(1-19: signal sequence) | Cys55Ala/Cys58Ala |
| 17 | **P5-A5 (CxxC, AxxA)** | pET15b (Novagen) | Nde I-BamH I | His<sub>6</sub> at N-term | 20-440 a.a.<br>(1-19: signal sequence) | Cys190Ala/Cys193Ala |
| 18 | **pON106 (P5-a0ab-cMyc-KEEL)** | pcDNA3.1<sup>+</sup> (Invitrogen) | HindIII-BspEI | Myc at C-term | 1-440 a.a.<br>(1-19: signal sequence) |  |
| 19 | **P5-A5-cMyc-KEEL** | pcDNA3.1<sup>+</sup> (Invitrogen) | HindIII-BspEI | Myc at C-term | 1-440 a.a.<br>(1-19: signal sequence) | V124A, L128A, V131A,<br>V135A, L139A |
| 20 | **P5-WT-3xcMyc-KEEL** | pcDNA3.1<sup>+</sup> (Invitrogen) | HindIII-BspEI | 3xMyc at C-term | 1-440 a.a.<br>(1-19: signal sequence) |  |
| 21 | **P5-A5-3xcMyc-KEEL** | pcDNA3.1<sup>+</sup> (Invitrogen) | HindIII-BspEI | 3xMyc at C-term | 1-440 a.a.<br>(1-19: signal sequence) | V124A, L128A, V131A,<br>V135A, L139A |
| 22 | **P5-WT-3xcMyc-KDEL** | pcDNA3.1<sup>+</sup> (Invitrogen) | HindIII-BspEI | 3xMyc at C-term | 1-440 a.a.<br>(1-19: signal sequence) |  |
| 23 | **P5-A5-3xcMyc-KDEL** | pcDNA3.1<sup>+</sup> (Invitrogen) | HindIII-BspEI | 3xMyc at C-term | 1-440 a.a.<br>(1-19: signal sequence) | V124A, L128A, V131A,<br>V135A, L139A |
| 24 | **P5-NQ-3xcMyc-KDEL** | pcDNA3.1<sup>+</sup> (Invitrogen) | HindIII-BspEI | 3xMyc at C-term | 1-440 a.a.<br>(1-19: signal sequence) | E422Q, D423N, D424N,<br>D426N, D429N, E431Q,<br>D433N, D434N |
| 25 | **IRE1α LD-GST** | pGEX6p-1<br>(GE Healthcare) | Out of multi-cloning site | GST at C-term | 24-446 a.a.<br>(1-23: signal sequence) |  |
| 26 | **IRE1α LD-GST-6xHis** | pGEX6p-1<br>(GE Healthcare) | Out of multi-cloning site | GST-6xHis<br>at C-term | 24-446 a.a.<br>(1-23: signal sequence) |  |

**Supplementary Table 2.**
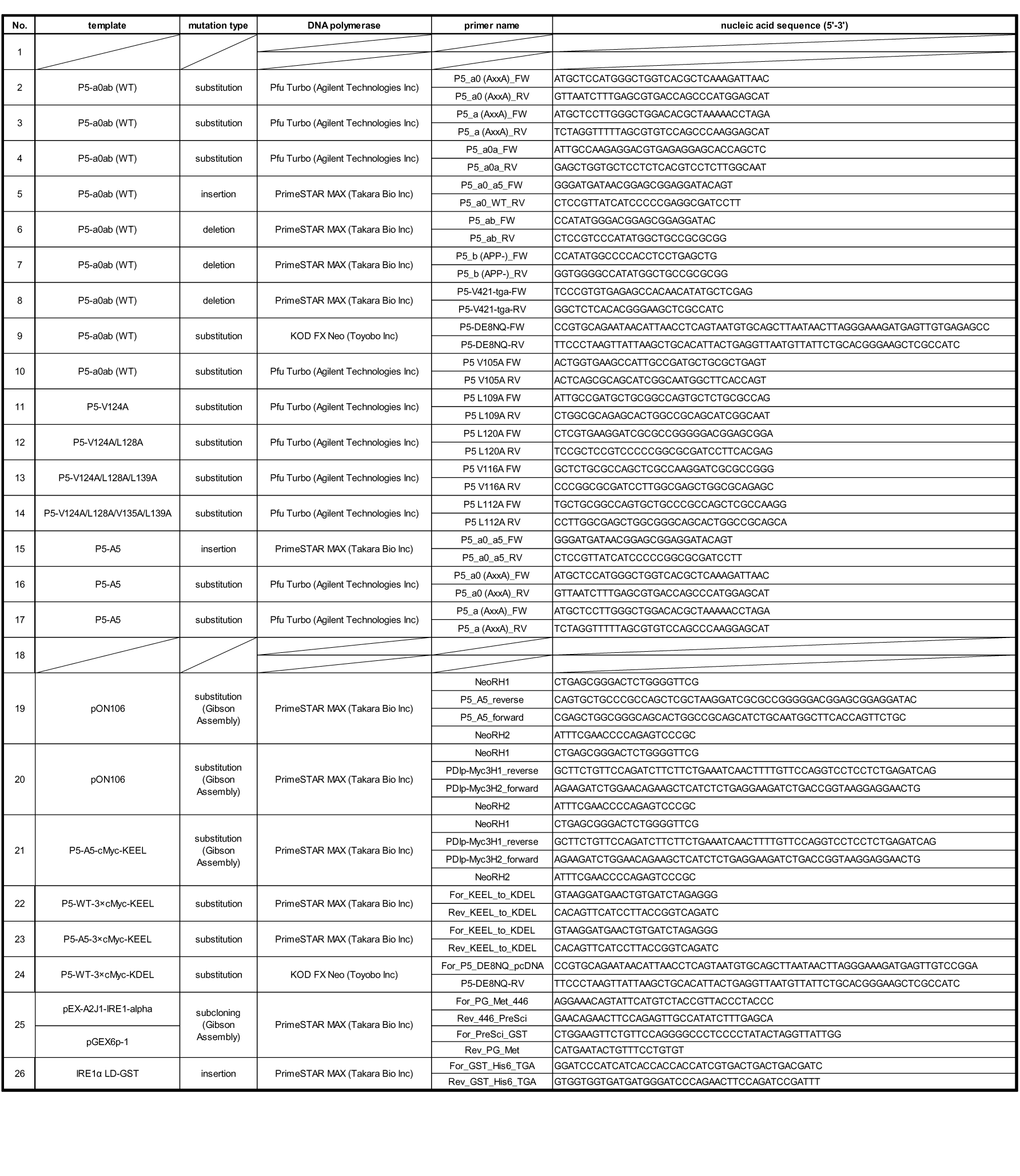
Primers for Cloning.

## METHODS

### Sample preparation

Human P5 used in this study was overexpressed and purified as described previously(Kojima et al., 2014). For preparation of reduced and oxidized P5, purified P5 was incubated in 50 mM TRIS-HCl buffer (pH 7.5) and 300 mM NaCl containing 1 mM dithiothreitol (DTT; reduced form) or 1 mM diamide (oxidized form) on ice for 10 min. After incubation, DTT or diamide was removed by gel filtration on a Superdex 200 10/300 Increase column (GE Healthcare, Chicago, USA). For isotopically-labeled samples used in nuclear magnetic resonance spectroscopy (NMR) studies, *Escherichia coli* cells harboring the plasmid for overexpression of human P5 or its mutants were grown in minimal (M9) medium at 37°C in the presence of ampicillin (100 mg/L). Protein samples for resonance assignment were expressed in M9 medium containing ^15^NH4Cl (1 g/L; CIL) and ^1^H^13^C-glucose (2 g/L; ISOTEC) in 99.9% ^2^H2O to prepare partially deuterated protein in which the protons derived from water were replaced with deuterium. Redox states of P5 thus prepared were confirmed by SDS-PAGE after modification with the thiol-alkylating agent maleimide-PEG-2K. Antibodies were from the following sources: anti-KDEL (Cat. No. M181-3; Medical & Biological Laboratories [MBL] Co., Ltd., Nagoya, Japan), anti-c-Myc (Cat. No. sc-40; Santa Cruz Biotechnology, Dallas, TX), anti-c-Myc (Cat. No. 562; MBL), anti-GAPDH (Cat. No. G9295; Sigma-Aldrich, St. Louis, MO), anti-PDI (Cat. No. ADI-SPA-891; Enzo Life Sciences, Farmingdale, USA), anti-HA agarose (Cat. No. #11815016001, Roche, Switzerland).

### Small-angle X-ray scattering (SAXS) measurements

Reduced and oxidized P5 and reduced P5 mutants (**a^0^a, ab, P5A5**) for SAXS measurements, and bovine serum albumin (BSA; molecular mass 66,400 Da; Sigma-Aldrich) as a reference for determination of the molecular mass were prepared as described previously(Akiyama, 2010). Oxidized P5 and BSA were dissolved in 20 mM phosphate buffer (pH 8.0) containing 150 mM NaCl and 5% glycerol. Reduced P5 and P5 mutants were dissolved in 20 mM phosphate buffer (pH 8.0) containing 150 mM NaCl, 5% glycerol, and 1 mM DTT. Small-angle X-ray scattering (SAXS) measurements were conducted at the SPring-8 RIKEN beamline BL45XU (Hyogo, Japan). For each sample, 20 diffraction images were collected using a PILATUS 3X 2M detector (DECTRIS) at an X-ray wavelength of 1.0 Å with a camera distance of 2.0 m and an exposure time of 1 s at 20.2°C. Data processing resulted in scattering patterns, *I*(*Q*), where *Q* = 4πsinθ/λ, 2θ is the scattering angle, and λ is the wavelength of the X-rays. The scattering profiles in the small-angle region were fitted under the Guinier approximation to the equation *I*(*Q*) = *I*(0)exp{-*Rg*^2^*Q*^2^/3}, where *I*(0) is the forward scattering intensity (*Q* = 0) and *R*g is the radius of gyration. The *I*(0) value is proportional to the averaged molecular mass. Pair-distribution functions, *P*(*r*), were calculated by indirect Fourier transformation using GNOM (Svergun, 1991).

### Plasmid construction

Plasmids and primers used in this study are listed in Supplementary Table 1 and 2. The mammalian expression plasmid containing 3×cMyc-tagged P5 and plasmid No. 18 (pON106, containing P5-cMyc-KEEL) were constructed previously(Fujimoto, Nakamura et al., 2018). Plasmid No. 19 (P5A5-cMyc-KEEL) was constructed by assembling two fragments amplified from pON106 using two primer pairs (NeoRH1 and P5_A5_reverse, P5_A5_forward and NeoRH2) with Gibson Assembly Master Mix (New England Biolabs, Ipswich, MA, USA). Plasmid No. 20 (P5-3×cMyc-KEEL) and No. 21 (P5-A5-3×cMyc-KEEL) were constructed by assembling two fragments amplified from pON106 with two primer pairs (NeoRH1 and PDIp-Myc3H1_reverse, PDIp-Myc3H1_forward and NeoRH2) and from P5-A5-cMyc-KEEL with two primer pairs (NeoRH1 and PDIp-Myc3H1_reverse, PDIp-Myc3H1_forward and NeoRH2) with Gibson Assembly Master Mix. Plasmid No. 22 (P5-3×cMyc-KDEL) and No. 23 (P5-A5-3×cMyc-KDEL) were generated by site-directed mutagenesis from P5-3×cMyc-KEEL and P5-A5-3×cMyc-KEEL, respectively, with primer pair For_KEEL_to_KDEL and Rev_KEEL_to_KDEL. Plasmid No. 24 (P5-NQ-3×cMyc-KDEL) was generated by site-directed mutagenesis from P5-WT-3×cMyc-KDEL using primer pair For_P5_DE8NQ_pcDNA and P5-DE8NQ-RV.

For the expression plasmid encoding the IRE1 luminal domain (LD), the full-length IRE1α gene was synthesized by Eurofins Genomics (Tokyo, Japan) according to the cDNA sequence reported in the NCBI database (GI:2081; https://www.uniprot.org/uniprot/O75460)] with codon optimization for expression in *E. coli*, and cloned into pEX-A2J1-IRE1-alpha. The appropriate DNA fragment of IRE1α LD was amplified by PCR from pEX-A2J1-IRE1-alpha using primer pair For_PG_Met_446 and Rev_446_PreSci. The pGEX-6P-1 vector (GE Healthcare, Uppsala, Sweden) was also amplified and modified by inverse PCR using primer pair For_PreSci_GST and Rev_PG_Met. For plasmid No. 25 (IRE1α LD-GST), the fragment was cloned into the modified pGEX-6P-1 vector using Gibson Assembly Master Mix, yielding a construct harboring the IRE1α LD gene fused to a C-terminal glutathione S-transferase (GST) tag separated by a PreScission Protease site. For plasmid No. 26 (IRE1α LD-GST-6×His), an additional 6×His-tag sequence was inserted by inverse PCR with primer pair For_GST_His6_TGA and Rev_GST_His6_TGA.

### Cell culture and transfection

HeLa Kyoto cells were kindly gifted from Dr. Kozo Tanaka (Tohoku University). HeLa Kyoto cells were cultured in Dulbecco’s modified Eagle’s medium (1.0 g/L glucose) containing L-Gln, sodium pyruvate (Nacalai Tesque, Kyoto, Japan), and 10% fetal bovine serum (FBS; Gibco). Plasmids were transfected with FuGENE HD Transfection Reagent (Promega).

### Immunoblotting

For assessment of BiP upregulation by P5 overexpression, HeLa Kyoto cells (5.0×10^4^ cells) were plated in a 6-well plate and transfected with plasmids after 24 h of incubation. In one well of a 6-well plate, 200 µL Opti-MEM, 1.2 µL FuGENE HD, and an optimized amount of plasmid DNA (Empty, 0.3 µg; P5WT, 0.4 µg; P5A5, 1.0 µg) were mixed. Transfected cells were incubated for an additional 40 h and harvested in SDS sample buffer containing 40 mM N-ethylmaleimide (NEM). Samples were separated on 7.5% SDS-PAGE gels, transferred to polyvinylidene fluoride (PVDF) membranes, and blotted with anti-KDEL (Cat No. M181-3; MBL, 1:10,000 in 1% MTBST: 50 mM Tris-HCl pH 7.6, 150 mM NaCl, 0.05% v/v Tween-20, 1% w/v skim milk), anti-c-Myc (Cat No. sc-40; Santa Cruz Biotechnology, 1:5000 in 1% MTBST), and anti-GAPDH (Cat No. G9295; Sigma-Aldrich, 1:7000 in 1% MTBST) antibodies. The band intensities of GRP94, BiP, and GAPDH were analyzed by a ChemiDoc Touch imaging system and Image Lab 5.2.1 (Bio-Rad). Statistical analyses were performed using one-way ANOVA and Tukey-Kramer tests.

### Immunofluorescence

To analyze the localization of exogenously expressed P5-3×cMyc-KDEL and P5-A5-3×c Myc-KDEL in HeLa Kyoto cells, cells were plated onto poly-L-lysine coated coverslips. Transfection was performed as described above for immunoblotting. At 36 h after transfection, cells were washed twice with phosphate-buffered saline (PBS) at room temperature (RT), and fixed with glyoxal solution (3% v/v glyoxal, 20% v/v ethanol, 128 mM acetic acid)(Richter, Revelo et al., 2018). Glyoxal (40% stock) was purchased from Sigma-Aldrich (#128465). Fixation was performed first for 30 min on ice, then for 30 min at RT. Fixed cells were washed four times with PBS at RT, permeabilized with PBS containing 0.1% Triton X-100 for 15 min, and blocked with PBS containing 2% FBS for 1 h at RT. Cells were then incubated overnight at 4°C with antibodies against c-Myc (Cat. No. 562; MBL, 1:1000) and PDI (Cat. No. ADI-SPA-891; Enzo Life Sciences, 1:1000) diluted in PBS containing 2% FBS. Following incubation, samples were washed three times with PBS at RT. Cells were incubated at 4°C for 4 h with CF488-conjugated anti-mouse IgG (1:2000) and CF568-conjugated anti-rabbit IgG (1:2000) antibodies (Biotium) diluted in PBS containing 2% FBS as secondary antibodies. Finally, nuclei were stained with 4′,6-diamidino-2-phenylindol (DAPI; 11034; Nacalai Tesque) and fluorescence images were obtained using a laser scanning confocal microscope (FV1000, Olympus) equipped with a UPLSAPO 60× silicon oil immersion objective lens (NA 1.30).

### Recombinant protein expression and purification of IRE1α LD

IRE1α LD was overexpressed in *E. coli* strain BL21 (DE3) and purified by optimized procedures established for disulfide-linked IRE1α (Extended Data Fig. 6). Expression of recombinant proteins was induced by adding 0.5 mM isopropyl-β-D-thiogalactopyranoside and culturing cells at 15°C overnight. Cells were disrupted using an NS1001L 2K homogenizer (Niro Soavi) in buffer containing 20 mM sodium phosphate (pH 7.2), 0.3 M NaCl, and 10% (w/v) glycerol. After clarification of the cell lysate by centrifugation (20,000×g for 20 min), the supernatant was purified by Glutathione Sepharose 4B column chromatography (GE Healthcare) followed by digestion with PreScission protease (GE Healthcare) and further purification by RESOURCE Q 6 mL column chromatography (GE Healthcare). Finally, samples were purified by Superdex 200 increase 10/300 GL column chromatography (GE Healthcare) with the column pre-equilibrated in buffer containing 25 mM HEPES (pH 7.2), 25 mM L-arginine, and 150 mM NaCl.

### Disulfide-linked IRE1α LD reduction assay

Purified disulfide disulfide-linked IRE1 LD (25 µM) was incubated with 250 µM TCEP with or without 0.5 µM P5WT or its mutants in 25 mM HEPES (pH 7.2), 25 mM L-arginine, and 150 mM NaCl at 30°C. The reactions were quenched with SDS-PAGE sample buffer containing 40 mM NEM at selected time points. The quenched samples were separated on 7.5% polyacrylamide gels with WIDE RANGE Gel Preparation Buffer (Nacalai Tesque) and stained with Coomassie Brilliant Blue G-250 (Nacalai Tesque). The band intensities of disulfide-linked IRE1α LD were analyzed by a ChemiDoc Touch imaging system and Image Lab 5.2.1 (Bio-Rad).

### BPTI refolding assay

Reduced and denatured BPTI (50 μM) was dissolved in 50 mM HEPES (pH 7.5) containing 150 mM NaCl, 1 mM GSH, 0.2 mM GSSG, and P5/PDI, as described previously(Kojima et al., 2014, Okumura, Kadokura et al., 2014, Okumura et al., 2019). All solutions used in this experiment were flushed with N2 gas, and the reactions were carried out in a sealed vial under an N2 atmosphere at 30°C. The reaction mixture (200 μL aliquots) was quenched with an equivalent volume of 1 M HCl at the indicated time points and separated by RP-HPLC on a TSKgel Protein C4-300 column (4.6 × 150 mm; Tosoh Bioscience) with monitoring at 229 nm. The identities of the resulting peaks were confirmed by MALDI-TOF/MS analysis as described above(Okumura, Saiki et al., 2011). Values are presented as means ± standard deviation (SD) from three independent experiments.

### Insulin reductase assay

Disulfide reductase activity was assessed by measuring the glutathione-dependent reduction of insulin according to a modified method from a previous study(Maegawa, Watanabe et al., 2017, Matsusaki, Okuda et al., 2016, Morjana & Gilbert, 1991). Recombinant proteins (1 µM) were incubated at 30°C in 80 µL of 50 mM HEPES buffer (pH 7.5) containing 150 mM NaCl, 10 mM glutathione (Sigma-Aldrich), 200 µM NADPH (Oriental Yeast Co., LTD.), 0.16 U of glutathione reductase (Sigma-Aldrich), and 30 µM bovine insulin (Sigma-Aldrich), and the absorbance was monitored at 340 nm with a U-3310 spectrophotometer (Hitachi High-Technologies, Tokyo, Japan). Relative insulin consumption rates were calculated in phases where the change in absorbance was stable. Statistical analysis was performed using one-way ANOVA and Tukey-Kramer tests.

### ITC measurements

Isothermal titration calorimetry (ITC) was carried out using a MicroCal VP-ITC instrument (Malvern Panalytical, United Kingdom) at 25 °C. Solutions of P5 or its variants (50 μM) and CaCl2 (10 mM) were prepared in 50 mM HEPES buffer (pH 7.5) containing 50 mM NaCl. All the samples were degassed for 3 min at 25 °C using the ThermoVac unit (Malvern Panalytical, United Kingdom) before ITC measurements. The solution of CaCl2 in the syringe was titrated into the solution containing P5 proteins in the cell with 28 injections at a constant interval of 600 s. The injection volume is 2 μl for the first injection and 10 μl for the remaining injections. The stirring speed of the syringe and the initial delay were set to 307 rpm and 600 s, respectively. Changes in the heat flow, i.e., ITC thermogram, were traced in real time with 10 μcal s^-1^ of reference power. The binding isotherms after the baseline correction and the subtraction of heat of dilution were fitted to a one-set of sites binding model incorporated in the MicroCal Origin software.

### CD measurement

Far-UV CD measurements were performed on a J-1500 spectrophotometer (JASCO, Tokyo, Japan) using a 0.1 cm path length quartz cell. Solutions of P5-WT and its variants were prepared at a concentration of 0.15 mg ml^-1^ in 20 mM sodium phosphate buffer (pH 7.4). Far-UV CD spectra of all P5 samples were acquired every 5 °C from 25 to 100 °C with a temperature increasing rate of 1 °C min^-1^. The temperature of the cell was controlled using RW-0525G Low Temperature Bath Circulator (Lab Companion, Daejeon, Republic of Korea). CD signals were converted to mean residue ellipticity, [*θ*] (degrees cm^2^ dmol^-1^)(Greenfield, 2006). The melting temperature (*T*m) and the enthalpy change (Δ*H*) for the heat denaturation of P5 and P5A5 were obtained by a regression analysis as described previously(Koepf, Petrassi et al., 1999).

### NMR measurements

For NMR experiments, isotopically labeled P5 **a^0^** and P5A5 **a^0^** were prepared in 20 mM sodium phosphate (pH 7.0), 100 mM NaCl, 5 mM DTT, and 10% ^2^H2O. ^15^N-labeled proteins were concentrated to 0.1 mM. For resonance assignment, ^13^C/^15^N-labeled proteins were concentrated to 0.3 mM. NMR spectra were recorded on a Bruker Avance Neo 800 MHz NMR spectrometer equipped with a cryogenic probe. Experiments were performed at 22°C. Spectra were processed using the NMRPipe program(Delaglio, Grzesiek et al., 1995), and data analysis was performed with Olivia (https://github.com/yokochi47/Olivia) or SPARKY . For resonance assignment, two-dimensional TROSY H(N)CA and H(NCO)CA, and three-dimensional TROSY HNCO spectra were recorded for ^13^C^15^N-labeled P5 **a^0^**. NMR signals were assigned using BMRB data as a reference (BMRB code: 11100).

### Gel shift assay for detection of disulfide-linked P5 oligomers under various redox conditions

P5 and its mutant proteins (2 μM) were incubated for 30 min at 30°C in degassed 50 mM HEPES (pH 7.5) buffer containing 0.2 mM GSSG and various concentrations of GSH (0.1–10 mM). After incubation, 20 mM NEM was added to reaction solutions. All samples were separated by non-reducing SDS-PAGE and stained with Coomassie Brilliant Blue (CBB). Band intensity was measured for monomeric and oligomeric states of P5 and its mutants using an LAS-3000 image reader (Fujifilm).

### Chaperone activity assay

Chaperone activities of P5 and its mutant proteins were measured using citrate synthase (CS) as a model substrate. To monitor the effect of P5 and its mutants on the thermal aggregation of CS, the substrate protein was diluted to a final concentration of 1 μM in buffer containing 40 mM HEPES-KOH (pH 7.5) and P5 (8 μM) in the presence or absence of 1 mM CaCl2. Protein aggregation was induced under agitation for 5 s at 43°C, and monitored at 350 nm using an SH-9000 microplate reader (Corona Electric Co., Ibaraki, Japan).

### Data availability

None of the data reported in this work have been deposited in public databases.

